# Evolution of multidrug-resistant IncC plasmids in the seventh pandemic *Vibrio cholerae* O1 El Tor lineage between 1979 and 2024

**DOI:** 10.64898/2026.05.05.723028

**Authors:** Rayane Rafei, Elisabeth Njamkepo, Marine Gaillard, Sahra Isse Mohamed, Anthony M. Smith, Nabila Benamrouche, Carlo Pazzani, Thandavarayan Ramamurthy, Fouad Dabboussi, Monzer Hamze, Marie-Laure Quilici, Caroline Rouard, François-Xavier Weill

**Affiliations:** Laboratoire Microbiologie Santé et Environnement (LMSE), Doctoral School of Sciences and Technology, Faculty of Public Health, Lebanese University, Tripoli, Lebanon; Institut Pasteur, Université Paris Cité, Unité des Bactéries pathogènes entériques, Centre National de Référence des Vibrions et du choléra, Paris, F-75015, France; Integrated Disease Surveillance & Response Unit, Ministry of Health & Human Services, Mogadishu, Somalia; Centre for Enteric Diseases, National Institute for Communicable Diseases, Division of the National Health Laboratory Service, Johannesburg, South Africa; Department of Medical Microbiology, School of Medicine, Faculty of Health Sciences, University of Pretoria, Pretoria, South Africa; Enterobacteria and Other Related Bacteria Laboratory, Pasteur Institute of Algeria, Algiers, Algeria; Faculty of Medicine, University of Health Sciences, Algiers, Algeria; University of Bari “A. Moro”, Department of Biosciences, Biotechnologies and Environment, Bari, Italy; Division of Bacteriology, ICMR-National Institute for Research in Bacterial Infection, Kolkata, West Bengal, India; Clinical Laboratory, Nini Hospital, Microbiology Department, Tripoli, Lebanon

**Keywords:** *Vibrio cholerae*, cholera, IncC plasmids, antimicrobial resistance genes, evolution

## Abstract

IncC plasmids played an important role in antimicrobial resistance development in the seventh pandemic *Vibrio cholerae* O1 El Tor (7PET) lineage during the 1970s and have recently begun to re-emerge. We investigated 28 complete IncC plasmids, including 17 newly sequenced genomes, from 7PET isolates collected across Asia, Africa, and the Caribbean between 1979 and 2024 and spanning the global 7PET phylogeny. Four additional IncC plasmids from other species were included, for a total of 32 IncC plasmids. IncC type 2 predominated among *V. cholerae* plasmids, and five novel core genome sequence types (cgSTs) were identified. Antimicrobial resistance genes (ARGs) occurred within diverse genetic structures inserted at specific sites in the conserved IncC backbone. Despite remarkable diversity in ARGs and associated genomic islands, the IncC backbone remained stable over five decades. These findings highlight the dynamic evolution and adaptability of IncC plasmids in 7PET under antimicrobial selection.

## INTRODUCTION

Cholera is a severe acute gastrointestinal illness caused by the two toxigenic serogroups of *Vibrio cholerae*: O1 and O139 ^1^. Cholera remains a global threat. The current, seventh pandemic began in 1961 and is estimated to cause 1.3 to 4.0 million cases and 21,000 to 143,000 deaths annually ^2^. This pandemic has included many major cholera outbreaks reported in different regions worldwide, including Yemen, Haiti, and Sudan, particularly in the wake of the breakdown of hygienic and sanitary conditions during natural disasters, wars, and massive human displacements ^3–5^. Genomic studies have revealed that the sustained propagation of the seventh cholera pandemic is due to a single lineage known as 7PET (seventh pandemic *V. cholerae* O1 El Tor) ^6–8^. This lineage first emerged in Indonesia and has repeatedly spread from the Bay of Bengal to different parts of the world, in at least three independent but temporally overlapping waves ^8^. Wave 1 emerged in the early 1960s, wave 2 in the 1980s, and wave 3 in the 1990s ^7,8^. Based on phylogenetic analysis, these waves formed three distinct genomic groups within the 7PET lineage, each with numerous sublineages ^7,8^. For instance, 11 different sublineages (AFR1, AFR3-AFR12) of the 7PET lineage were introduced into Africa from South Asia between 1970 and 2014 ^7^. Other sublineages (AFR13 to AFR15) responsible for cholera outbreaks in Africa over the next decade have since been described ^9–11^.

One major concern during the cholera outbreaks of this pandemic is the emergence of multidrug-resistant (MDR) strains, which may compromise the efficacy of antibiotic treatments recommended alongside fluid and electrolyte replacement therapy ^12–14^. Conjugative plasmids from the IncA and IncC incompatibility groups (historically referred to as the IncA/C group) ^15^ are one of the major drivers of antimicrobial resistance (AMR) in Gram-negative bacteria, including *V. cholerae,* due to their broad host range and contribution to the dissemination of AMR genes (ARGs), including some encoding clinically important extended-spectrum beta-lactamases (ESBLs) and carbapenemases. Comparative genomic analyses have distinguished two major groups of A/C plasmids, A/C1 (IncA) and A/C2 (IncC) ^16,17^. Subsequent analyses further classified IncC plasmids into IncC type 1 (IncC1) and IncC type 2 (IncC2), based on four genomic segments: i1, i2, Region 1 (R1), and Region 2 (R2) ^16^. The i1 and i2 segments are approximately 600 bp in IncC2, compared with approximately 200 bp in IncC1, owing to insertions of ∼400 bp. R1 contains a large ORF between *traA* and *dsbC* (*orf1832* in IncC1 and *orf1847* in IncC2), whereas R2 comprises *rhs1* in IncC1 or *rhs2* in IncC2 and a downstream variable region ^16^. AMR genes are generally clustered within resistance islands on IncC plasmids ^16^, notably ARI-B, derived from the 15,460-bp genomic island GI*sul2* and carrying the sulfonamide resistance gene *sul2*, and ARI-A, a complex AMR-associated transposon.

In the 1970s and 1980s, MDR plasmids encoding resistance to several antibiotics, such as tetracycline, trimethoprim-sulfamethoxazole, and aminoglycosides, were described in 7PET isolates, and some of these plasmids were associated with outbreaks ^18–21^. These MDR plasmids belonged to the IncC group, suggesting that IncC plasmids may be particularly well adapted to *V. cholerae* ^18^. Consistent with this hypothesis, an earlier study showed that plasmids from most other Inc groups were unstable in *V. cholerae*, whereas IncC plasmids and the R391 element were stably maintained over 200–400 generations ^22^. R391, originally identified in *Proteus rettgeri* (now *Providencia rettgeri*) and initially classified as an IncJ plasmid, is now recognized as an integrative and conjugative element (ICE) of the SXT/R391 family ^23,24^. However, IncC plasmids, which were frequently detected towards the end of wave 1 of the seventh cholera pandemic, subsequently became rare thereafter ^7^. This decline was proposed to result from the acquisition by 7PET isolates of SXT/R391 ICEs, particularly ICE*Vch*Ind5, which carry ARGs and have been present in 7PET strains for the past 25 years. These ICEs became predominant in wave 2 and 3 strains, whereas wave 1 strains lacked SXT/R391 ICEs ^7,25^. Indeed, an apparent negative association between SXT/R391 ICEs and IncC plasmids was observed in 7PET strains: the vast majority of strains carrying SXT/R391 ICEs lacked IncC plasmids ^26^, whereas coexistence of both elements has been reported in the recent AFR13 sublineage. The apparent negative association between SXT/R391 ICEs and IncC plasmids has been proposed to result from functional interference between these two related MDR elements, although the underlying mechanism remains unknown ^7,24^. In addition, two chromosomally encoded defense systems (DdmABC and DdmDE) have been proposed to contribute to the low prevalence of plasmids, including IncC plasmids, within the 7PET lineage ^27,28^. The *ddmABC* and *ddmDE* loci are located within the chromosomal *Vibrio* seventh pandemic island II (VSP-II) and *Vibrio* pathogenicity island 2 (VPI-2), respectively. Both systems have been shown to decrease the acquisition of large plasmids in 7PET strains. In the absence of selection, DmdABC may increase the burden imposed by large plasmids (including IncC), resulting in a higher fitness cost that favors plasmid-free cells ^28,29^. IncC plasmids are nevertheless increasingly being detected in 7PET *V. cholerae* strains ^12,30,31^, and are stably maintained in successful sublineages, such as AFR13 ^32^. AFR13 strains have acquired distinct IncC plasmids on multiple occasions, including pYA00120881 and pCNRVC190243. pYA00120881 was identified in Zimbabwe during a cholera outbreak caused by a highly drug-resistant strain ^31^, whereas pCNRVC190243 was found in AFR13 isolates associated with the dissemination of high-level drug resistance across several countries in the Middle East and East Africa ^12,32^. The coexistence of IncC plasmids and ICE*Vch*Ind5 in AFR13 isolates has been proposed to result from a 10-kb deletion in the SXT/R391 ICE, potentially affecting the putative mechanism underlying their negative association ^12^. In addition to their role as a vehicle for ARGs, IncC plasmids can also mobilize specific chromosomal genomic islands, including *Salmonella* Genomic Island 1 (SGI1) and related elements, as well as MGI*Vch*Hai6 ^33^.

Most studies on *V. cholerae* plasmids have focused on the characterization of individual plasmids carried by outbreak isolates ^12,30–32^. Moreover, short-read sequencing approaches such as Illumina sequencing can be insufficient to resolve complex plasmid structures and AMR islands, leading to misassemblies and incomplete characterization. Here, we analyzed 28 complete IncC plasmids from 7PET strains with diverse spatiotemporal origins, spanning waves 1–3, multiple phylogenetic lineages, and nearly five decades of sampling. We further included four IncC plasmids and one IncA plasmid from other bacterial species, yielding a total of 33 A/C plasmids. After defining the conserved IncC plasmid backbone, we characterized the diversity and genomic organization of ARGs and associated AMR islands inserted in this backbone. Finally, we investigated the temporal evolution of IncC plasmids and their AMR islands across bacterial species and 7PET sublineages.

## METHODS

### Ethical statement

This study used only bacterial isolates and limited, non-identifiable metadata (isolation date and country of infection) collected through routine cholera surveillance in accordance with local regulations. Therefore, informed consent and ethics committee approval were not required.

### Global phylogenetic analysis of 7PET *V. cholerae* isolates

To represent the global phylogenetic context of the 7PET *V. cholerae* isolates, we used the most up-to-date and comprehensive 7PET genomic dataset available from Rouard et al. ^32^, comprising 1,523 genomes from the historical Pasteur collection and other published studies. This dataset encompasses the three pandemic waves and the major 7PET sublineages. We additionally included seven genomes for which plasmid sequences were available online or were generated in this study, resulting in a final dataset of 1,530 genomes. The paired-end reads and draft or assembled genomes of 1,530 7PET isolates (Supplementary Data 1) were mapped onto the reference genome of *V. cholerae* O1 El Tor N16961, also known as A19 (GenBank accession numbers LT907989 and LT907990), with Snippy version 4.6.0/BWA v.0.7.17 (https://github.com/tseemann/snippy). Single-nucleotide variants (SNVs) were called with Snippy v.4.6.0/Freebayes v.1.3.2 (https://github.com/tseemann/snippy) under the following constraints: mapping quality of 60, minimum base quality of 13, minimum read coverage of 4, and a read concordance of 75% at the locus concerned for a variant to be reported. An alignment of core genome SNVs was produced in Snippy for phylogeny inference.

Repetitive (insertion sequences and the TLC-RS1-CTX region, consisting of Toxin-Linked Cryptic element, RS1 satellite phage, and CTX cholera toxin phage) and recombinogenic (*Vibrio* seventh pandemic island, VSP-II) regions in the alignment were masked ^7^. Other putative recombinogenic regions were identified and masked with Gubbins ^34^ v.3.2.0. A maximum likelihood (ML) phylogenetic tree was built from an alignment of 10,978 chromosomal SNVs, with RAxML ^35^ v.8.2.12, under the GTR model with 200 bootstraps. This global tree was rooted on the A6 genome ^7^, the earliest and most ancestral 7PET isolate, collected in Indonesia in 1957, and visualized with iTOL ^36^ v.7.5 (https://itol.embl.de).

### Bacterial isolates

We studied 17 MDR seventh pandemic *V. cholerae* O1 El Tor isolates from the strain collection of the French National Reference Center for Vibrios and Cholera at the *Institut Pasteur*, Paris, France. Each isolate carried an IncC plasmid, and these isolates were broadly distributed across the 7PET *V. cholerae* phylogeny (Tables 1 and 2).

**Table 1:**
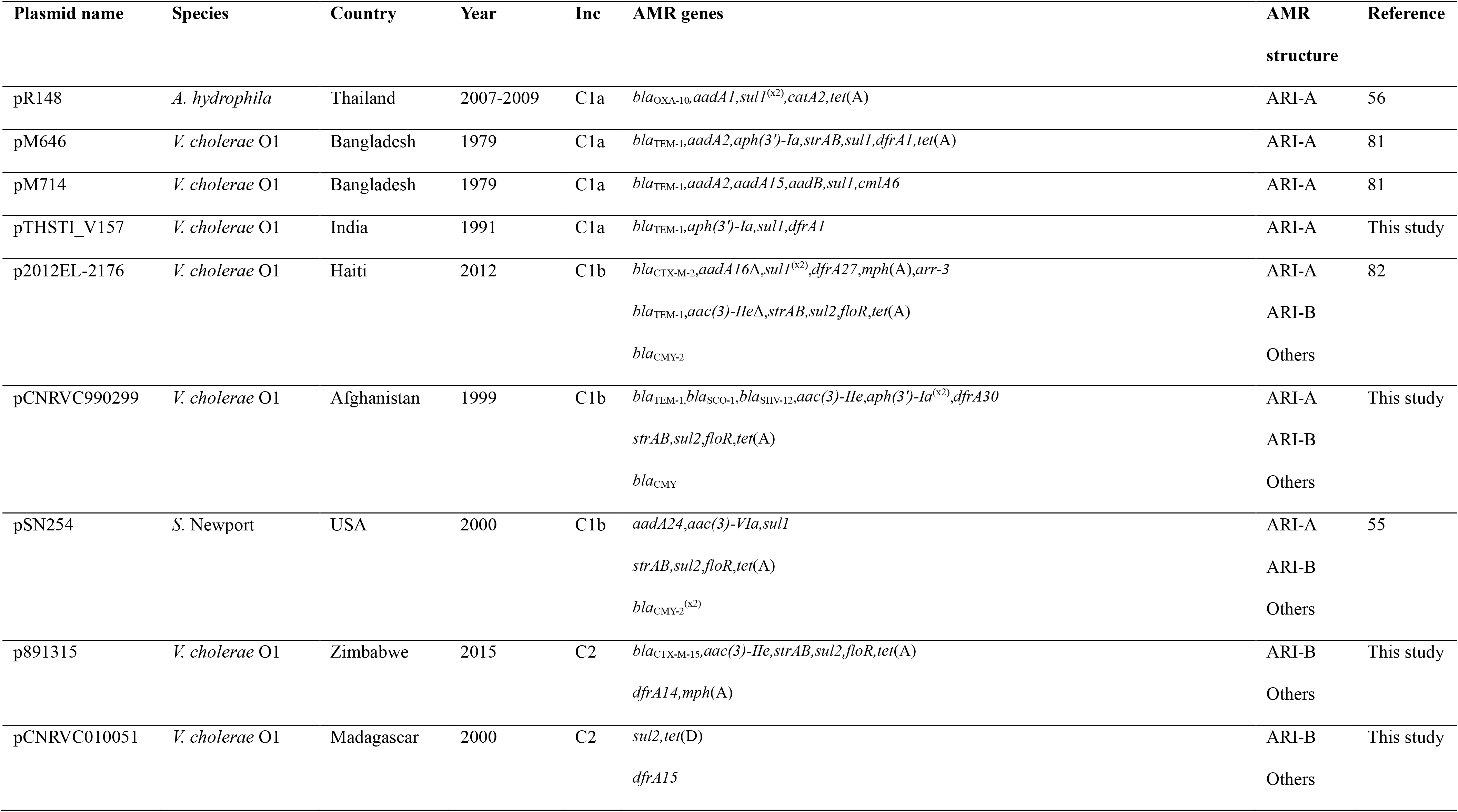
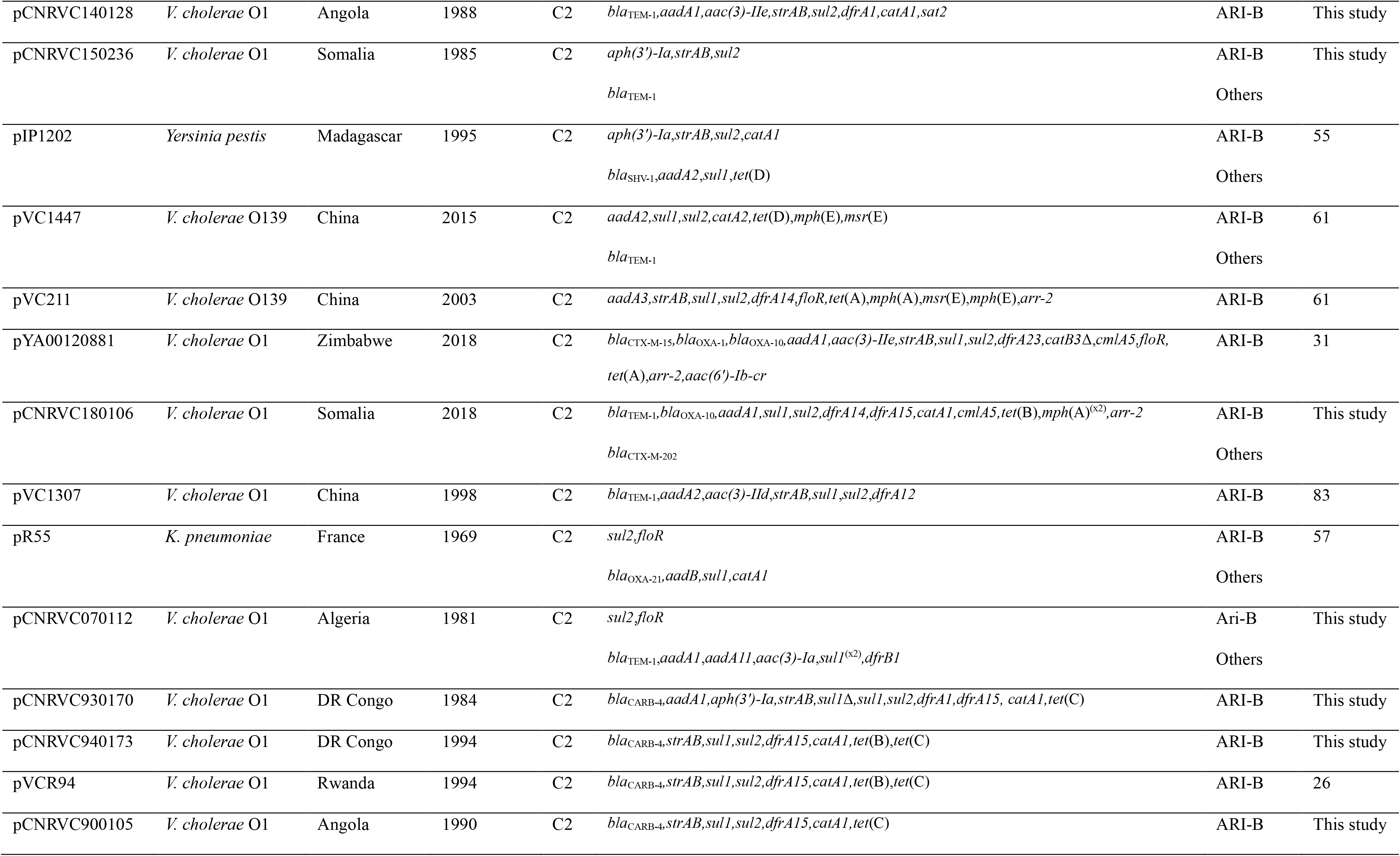
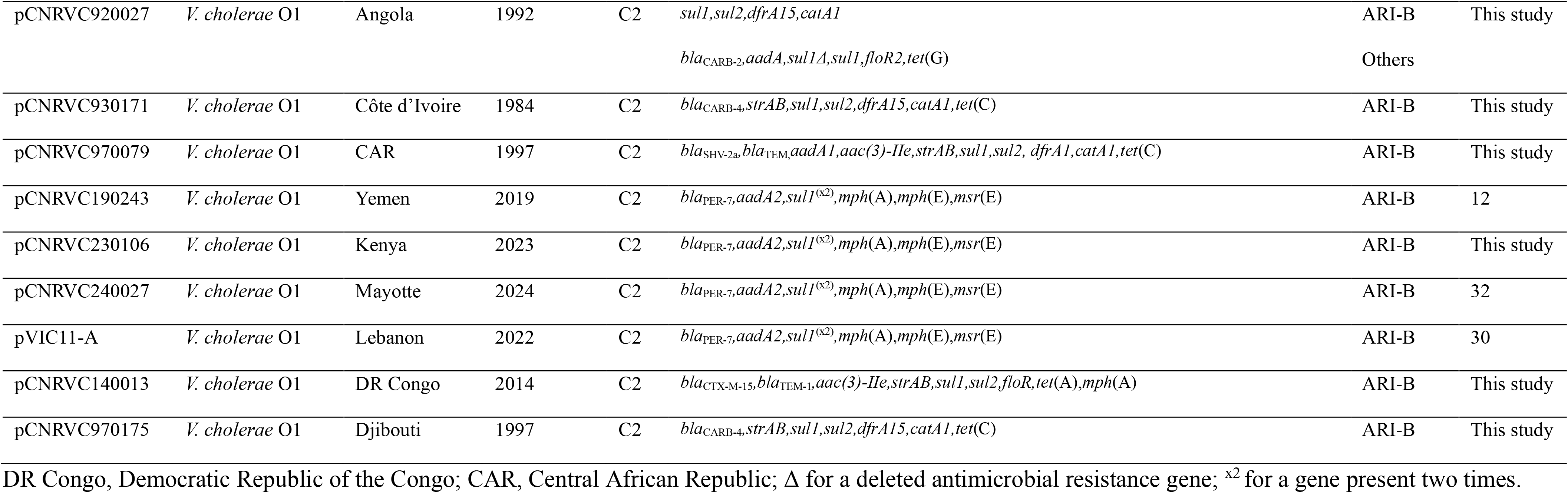
Key characteristics of the 32 IncC plasmids studied.

**Table 2:** Distribution of the studied plasmids according to introduction event in Africa and wave.

| Wave | Introduction event in Africa | Plasmid |
| --- | --- | --- |
| 1 | AFR1 | pCNRVC070112, pCNRVC140128, pCNRVC900105, pCNRVC920027, pCNRVC930171 |
|  | AFR5 | pCNRVC930170, pCNRVC940173, pVCR94*, pCNRVC150236 |
|  | NA | pM646, pM714 |
| 2 | AFR6 | pCNRVC970175 |
|  | AFR7 | pCNRVC970079 |
|  | NA | pTHSTI_V157, pVC1307, pVC1447, pVC211 |
| 3 | AFR10 | pCNRVC010051, pCNRVC140013 |
|  | AFR11 | p891315 |
|  | AFR13 | pYA00120881, pCNRVC180106, pCNRVC190243, pCNRVC230106, pCNRVC240027, pVIC11-A |
|  | NA | p2012EL-2176, pCNRVC990299 |

\*pVCR94 probably belonged to wave 1 and AFR5; plasmids, for which introduction into Africa is not applicable (NA).

### Antimicrobial drug susceptibility testing

Antimicrobial drug susceptibility testing (AST) data were already available for isolate 891315 ^31^. AST was performed on 16 *V. cholerae* O1 El Tor and two *E. coli* transconjugants from CNRVC180106 and CNRVC140013 by the microdilution method (Sensititre^TM^ Thermo Fisher Scientific, Cleveland, OH, USA) on two custom plates (FRVIB2 and FRVIB3). The EUCAST criteria for the interpretation of antibiotic susceptibility tests in *Vibrio spp*. (v. 16.0) were used when available (i.e., for cefotaxime (CTX), ceftazidime (CAZ), meropenem (MEM), ciprofloxacin (CIP), trimethoprim-sulfamethoxazole (SXT), azithromycin (AZI), doxycycline (DOX)) ^37^. For ampicillin (AMP), gentamicin (GEN), chloramphenicol (CHL), and sulfamethoxazole (SMX), the Clinical and Laboratory Standards Institute (CLSI) interpretation criteria for *Vibrio spp*. were used ^38^. For imipenem (IPM), trimethoprim (TMP), nitrofurantoin (NIT), and colistin (COL), the EUCAST interpretation criteria for Enterobacterales were used ^37^. For nalidixic acid (NAL), tetracycline (TET), and minocycline (MIN), the CLSI interpretation criteria for Enterobacterales were used ^39^. *Escherichia coli* ATCC 25922 and *Staphylococcus aureus* ATCC 29213 were used as internal quality controls.

### Long-read sequencing

We sequenced 17 MDR seventh pandemic *V. cholerae* O1 El Tor isolates carrying IncC plasmids (Supplementary Data 2). Eight of these isolates were from wave 1 (five AFR1 sublineage, three AFR5), three from wave 2 (one AFR6, one AFR7, and one from an Asian sublineage not formally designated), and six from wave 3 (two AFR10, two AFR13, one AFR11, and one from an Asian sublineage not formally designated). Total DNA was extracted from all isolates using MaXtract High Density phase-lock tubes (Qiagen, Germany), except for CNRVC230106 and 891315, where the Genomic-tip 100/G kit (Qiagen) and the QIAamp DNA Mini Kit (Qiagen) were used, respectively. The extracted DNA was sequenced on PacBio RS platforms (Pacific Biosciences, Menlo Park, CA, USA), except for CNRVC070112 and CNRVC230106, which were sequenced with Oxford Nanopore Technologies (Oxford, UK) using R9.4.1 flow cells with the Ligation Sequencing Kit (SQK-LSK109) and Native Barcoding Expansion (EXP-NBD104) (Supplementary Data 3). Long reads were assembled using Unicycler ^40^ v.0.4.8 in hybrid mode with Illumina reads, or using the Hierarchical Genome Assembly Process v.3 (HGAP3; Pacific Biosciences), followed by error correction through mapping of short reads, as described in Supplementary Data 3.

### Plasmid sequence analysis, typing, and phylogenetics

PlasmidFinder ^41^ v.2.1.1 was used to scan reads or assemblies of the 1,530 *V. cholerae* isolates for the presence of potential replicons. The 33 A/C plasmid sequences studied in this study, including the 17 new sequences, were annotated with Prokka ^42^ v.1.14.5. We screened the sequences for ARGs with ResFinder ^43^ v.4.3.3 and AMRFinderPlus ^44^ v.4.2.7. In cases of discordance, a final annotation was selected based on manual review; the results from both tools with the final assignment are provided in Supplementary Data 4. Insertion sequences (ISs) were identified with ISFinder ^45^. Transposon sequences were identified according to the TnCentral database ^46^. Integrons were identified by two approaches: with Integron Finder ^47^ v.2.0.5 or by manual BLAST analysis of the integrases and 5′ conserved and 3′ conserved regions of class 1 integrons. Integrons were annotated according to the Integrall database ^48^. The presence of potential CRISPR sequences was investigated with the online tool CRISPRCasFinder (https://crisprcas.i2bc.paris-saclay.fr/CrisprCasFinder/Index).

Plasmids were classified into IncA and IncC based on their replicons identified using PlasmidFinder ^41^ and into plasmid taxonomic units using COPLA ^49^. To classify plasmids into IncC1 and IncC2 groups, sequences of four segments (R1, R2, i1, i2) identified by Ambrose et al. ^16^ were retrieved from pR148 (IncC1, GenBank accession JX141473) and pR55 (IncC2, JQ010984) (Supplementary Data 5). These sequences were used to screen plasmids via BLASTn. IncC1 plasmids were further divided into IncC type 1a and 1b based on the presence of a high SNP density patch in the type 1a region, also identified using BLASTn. This region was extracted from pR148 as defined by Ambrose et al. ^16^. IncA and IncC plasmids were also attributed a sequence type (ST) and core genome ST (cgST) with the IncA/C plasmid multilocus sequence typing (PMLST) and IncA/C core genome plasmid multilocus sequence typing (cgPMLST) schemes available from PubMLST (https://pubmlst.org/organisms/plasmid-mlst), which are based on 4 and 28 core genes, respectively ^50^. New alleles and STs were also submitted to the database.

The different IncC plasmids were then compared to the pR55 backbone to identify the various modifications to the backbone that had occurred, including insertions containing ARGs. AMR islands that had already been defined were named as described by Ambrose et al. ^16^. BRIG ^51^ v.0.95 (https://sourceforge.net/projects/brig/) was used to visualize and compare plasmids.

For the IncC plasmid phylogeny, the 32 IncC backbones were aligned with Mafft ^52^ v.7.525 according to the FFT-NS-2 strategy. SNVs were extracted from the backbone plasmid alignment with SNP-sites ^53^ v.2.5.1. A maximum likelihood phylogenetic tree was inferred with RAxML ^54^ v.8.2.12, using GTRGAMMAI as the evolution model, with 1000 bootstrap replicates.

### Bacterial conjugation assays

Resistance transfer experiments were performed on solid media with *E. coli* K-12 J5 resistant to sodium azide as the recipient. Ten *V. cholerae* O1 El Tor isolates were selected as donor strains to maximize geographic, temporal, and phylogenetic representation across the three 7PET waves and their sublineages (Supplementary Data 6). All ten donor isolates were resistant to ampicillin due to the presence of various *bla* genes carried by the studied IncC plasmids (Table 1, Supplementary Data 6). Conjugation assays were performed by mixing equal volumes (100 µL) of six-hour cultures of the donor and recipient strains. The suspension was applied to a membrane filter on Luria Bertani (LB) agar (Difco, Becton Dickinson, Le Pont de Claix, France) and incubated at 37°C overnight. Cells were collected in 5 mL of saline and were serially diluted before plating. Transconjugants were selected on Drigalski agar (Bio-Rad, Marnes-la-Coquette, France) supplemented with sodium azide (200 mg/L) and ampicillin (100 mg/L). Three *E. coli* transconjugants were arbitrarily selected in each experiment.

### Data availability

All the data generated or analyzed during this study are included in the published article and its supplementary information files. The accession numbers for the plasmids sequenced in this study and for the publicly available data used in this study are provided in Supplementary Data 2. The genome of CNRVC180106 has been deposited in the European Nucleotide Archive (ENA) under BioProject accession number PRJEB80488 and run accession number ERR16793724.

## RESULTS AND DISCUSSION

In this study, we analyzed 32 IncC plasmids, including 28 IncC plasmids from 7PET *V. cholerae* isolates and four additional well-characterized IncC plasmids from *Yersinia pestis* ^55^, *Salmonella enterica* serotype Newport ^55^, *Aeromonas hydrophila* ^56^, and *Klebsiella pneumoniae*^57^. These included some of the first IncC plasmids to be sequenced (Table 1, Supplementary Data 2). The 28 IncC plasmids from 7PET were derived from 17 newly sequenced and 11 previously published isolates obtained from human clinical samples between 1979 and 2024 in 11 African countries, Mayotte (a French overseas department in the Indian Ocean, off the coast of East Africa), six Asian countries, and Haiti. These plasmids were recovered from 7PET isolates representing the three pandemic waves and seven sublineages distributed across the 7PET phylogeny (Fig. 1). Most 7PET isolates were multidrug-resistant, exhibiting phenotypic resistance to three to twelve antibiotics (Supplementary Data 2). We also included pRA1 from *A. hydrophila* ^58^, an IncA plasmid historically classified within the A/C incompatibility group. The 32 IncC plasmids and pRA1 were all classified by COPLA within PTU-C (Supplementary Data 2). Although sampling was uneven across time and geography, particularly because of the limited number of historical Asian isolates, this dataset represents a comprehensive collection of IncC plasmids from 7PET *V. cholerae* and enables comparison of their backbone and antimicrobial resistance architectures across diverse temporal and phylogenetic contexts.

**Figure 1:**
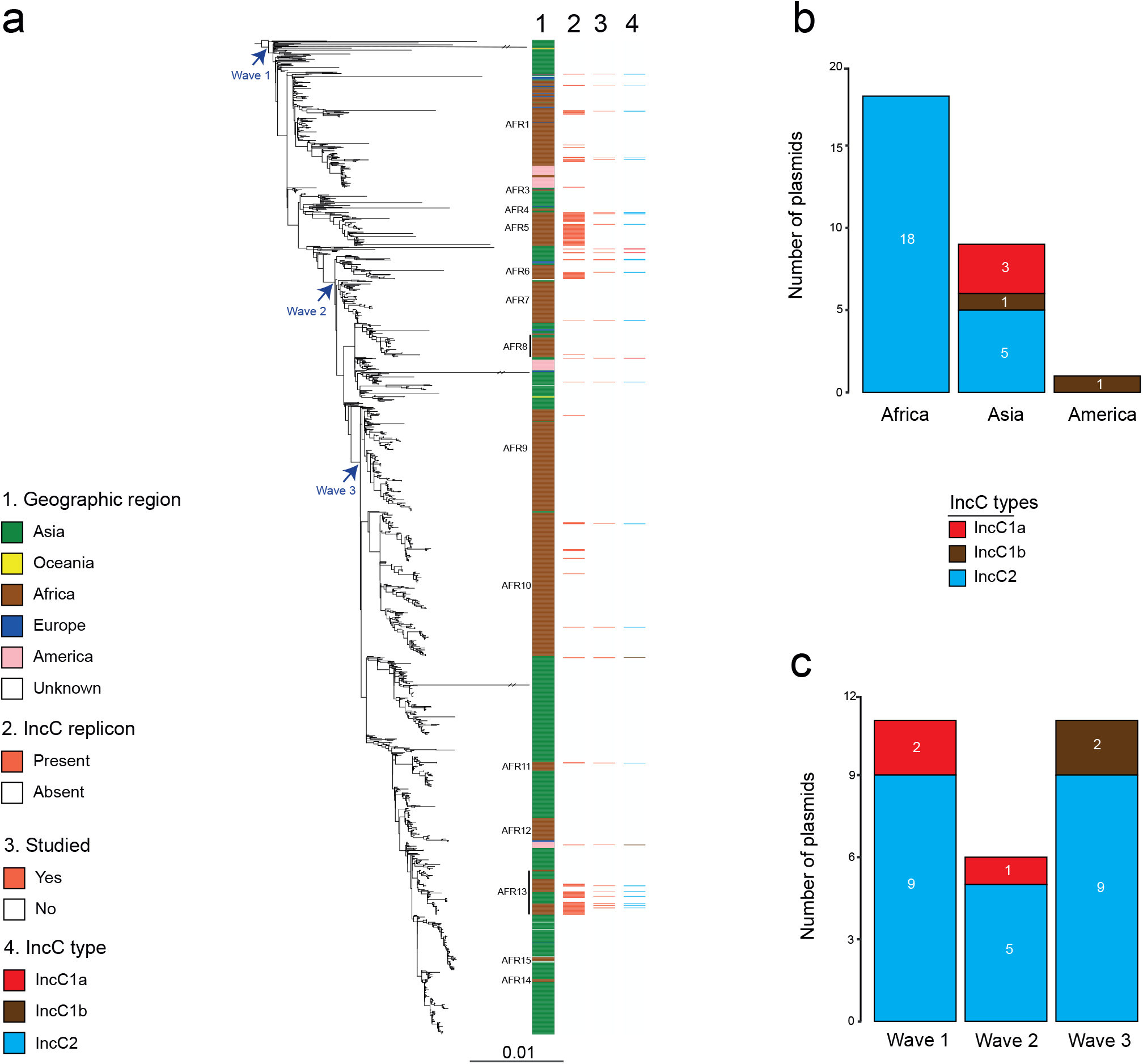
Maximum-likelihood phylogeny of 1,530 seventh pandemic *Vibrio cholerae* O1 El Tor isolates and distribution of IncC plasmids by geographic region and genomic wave, stratified by IncC plasmid type. **(a)** A6 was used as the outgroup. Blue arrows indicate the positions of the three genomic waves on the phylogenetic tree. Double slashes indicate an artificial shortening of the branches to enhance visualization. The scale bar indicates the number of nucleotide substitutions per variable site. The color key indicates the color scheme used in the four columns shown. The first column shows the geographic origin of the isolates, with African sublineages (AFR1 and AFR3–AFR15) shown on the left. The second column indicates the presence (red) or absence (white) of the IncC replicon, as determined by PlasmidFinder analysis of short-read assemblies from the 1,530 isolates. The red color in the third column identifies the plasmids studied here. The fourth column shows the classification of these plasmids into IncC types 2, 1a, and 1b, according to the colors used. The genome of the *V. cholerae* O1 El Tor isolate originally carrying pVCR94 (strain F1939, isolated in Rwanda in 1994) is not publicly available and was therefore not included in this phylogenetic analysis. **(b)** Distribution of the IncC plasmids included in this study across the three geographic regions represented in the dataset (Africa, Asia, and the Americas). Plasmids are colored according to their IncC type (IncC2, IncC1a, and IncC1b), as indicated in the color key. The number of plasmids belonging to each IncC type within each geographic region is indicated in white. **(c)** Distribution of the IncC plasmids across the three genomic waves of the seventh pandemic *V. cholerae* O1 El Tor lineage (Wave 1, Wave 2, and Wave 3). Plasmids are colored according to their IncC type (IncC2, IncC1a, and IncC1b), as indicated in the color key. The number of plasmids belonging to each IncC type within each genomic wave is indicated in white.

### IncC type 2 predominated among the 7PET *V. cholerae* plasmids studied

All 17 newly sequenced plasmids from 7PET *V. cholerae* were IncC plasmids, comprising 15 IncC2, one IncC1a, and one IncC1b plasmid (Supplementary Data 2 and 5). Overall, IncC2 predominated in our dataset, accounting for 25 of the 32 IncC plasmids, including 23 of the 28 plasmids from 7PET *V. cholerae* (Figs. 1B, 1C). The remaining seven plasmids were IncC1, comprising four IncC1a and three IncC1b plasmids. Among the IncC plasmids from 7PET *V. cholerae*, the two IncC types also showed distinct geographic distributions: four of the five IncC1 plasmids were associated with South and Southeast Asia, whereas IncC2 plasmids were predominantly recovered from Africa, with additional representatives from China and the Middle East. This pattern may reflect differences in the circulation or ecological persistence of the two IncC types in 7PET *V. cholerae*, although the limited number of IncC1 plasmids and uneven sampling warrant caution.

The IncC1 and IncC2 classification scheme proposed by Ambrose et al. ^16^, based on four backbone segments (i1, i2, R1, and R2), was largely applicable to the plasmids in our dataset, with some exceptions affecting the i1 segment. The larger i1 segment containing an approximately 400-bp insertion, previously associated with IncC2 plasmids, was detected in only six of the 25 IncC2 plasmids studied (Supplementary Data 5). Conversely, the shorter i1 segment generally found in IncC1 plasmids was absent from three IncC1 plasmids, all of which were IncC1b (Supplementary Data 5). The R2 region characteristic of IncC2 was conserved in all IncC2 plasmids except pCNRVC180106, which displayed a distinct terminal structure (Supplementary Data 5). In the IncC2 and IncC1b plasmids lacking the expected i1 segment, the deletion extended beyond i1 into an adjacent portion of the backbone, rather than affecting i1 alone, suggesting that these structural variants may have resulted from recombination events (Fig. 2). These deletions add further structural diversity to the otherwise conserved IncC backbone and may provide useful evolutionary and epidemiological markers.

**Figure 2:**
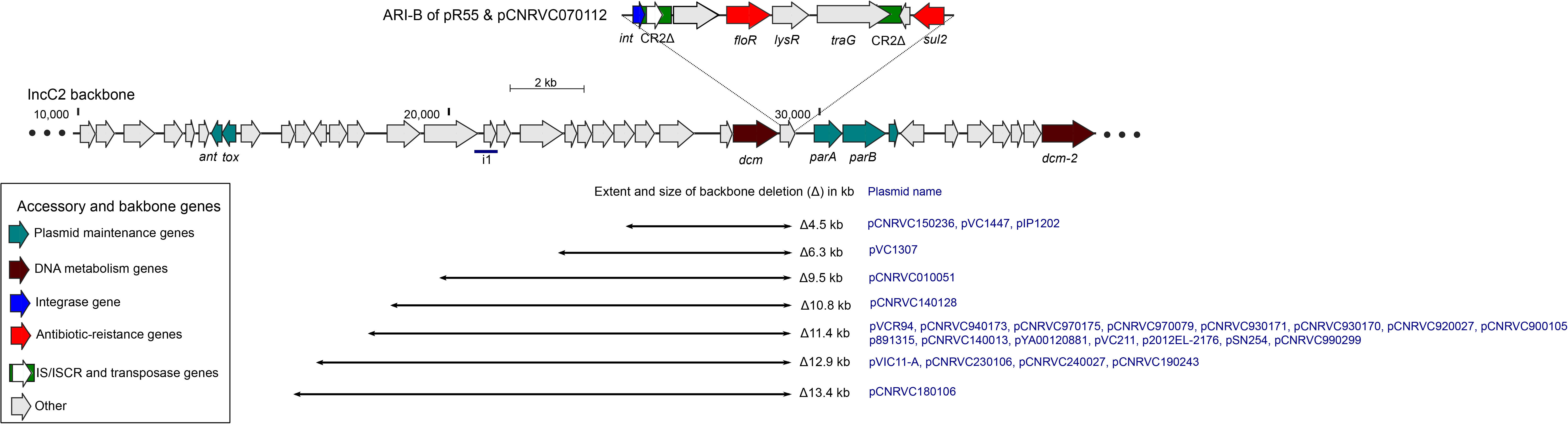
ARI-B-associated deletions in the backbone. The diagram shows a portion of the backbones of pR55 and pCNRVC070112, with ARI-B indicated above. The lines below represent deletions in different plasmids, with plasmid names shown in blue and deletion sizes indicated in kb to the right of each line. The length of each line is proportional to the size of the corresponding deletion. The line labeled “i1” below the backbone indicates the position of the i1 region.

In total, four A/C sequence types (STs) were identified among the 32 IncC plasmids studied and pRA1 (IncA), with ST3 predominating (28/33) and one novel ST (ST15) identified. These STs were further subdivided into 13 cgSTs, including five novel cgSTs, with ST3 showing the greatest diversity (10 cgSTs) and cgST3.3 being the most prevalent subtype (Supplementary Data 7). The backbone-based phylogeny separated the 32 IncC plasmids according to IncC type and was largely concordant with the cgST classification (Fig. 3). In our dataset, cgSTs were specific to individual IncC types or subtypes, with no cgST shared between IncC1a, IncC1b, and IncC2 plasmids. However, several cgSTs within IncC2, including cgST3.3, cgST3.25, cgST3.26, cgST3.5, cgST3.11, cgST3.14, and cgST3.27, were not clearly resolved by the backbone phylogeny. The difference between cgST3.14, represented by the previously published pR55 sequence, and cgST3.11, represented by pCNRVC070112, was due solely to an indel at the A002 locus of the cgPMLST scheme, within a homopolymeric region, which may reflect a sequencing or assembly artefact in the 454-generated pR55 sequence. Because the current cgPMLST scheme was developed using both IncA and IncC plasmids ^50^, an IncC-specific scheme may provide greater resolution of IncC plasmid diversity. Within IncC2, pCNRVC180106, recovered from Somalia in 2018, pCNRVC970175 from Djibouti in 1997, and the group comprising pCNRVC930170, pVCR94 ^26^, and pCNRVC940173 showed greater backbone divergence than the remaining IncC2 plasmids, consistent with the pairwise SNV analysis (Supplementary Data 8).

**Figure 3:**
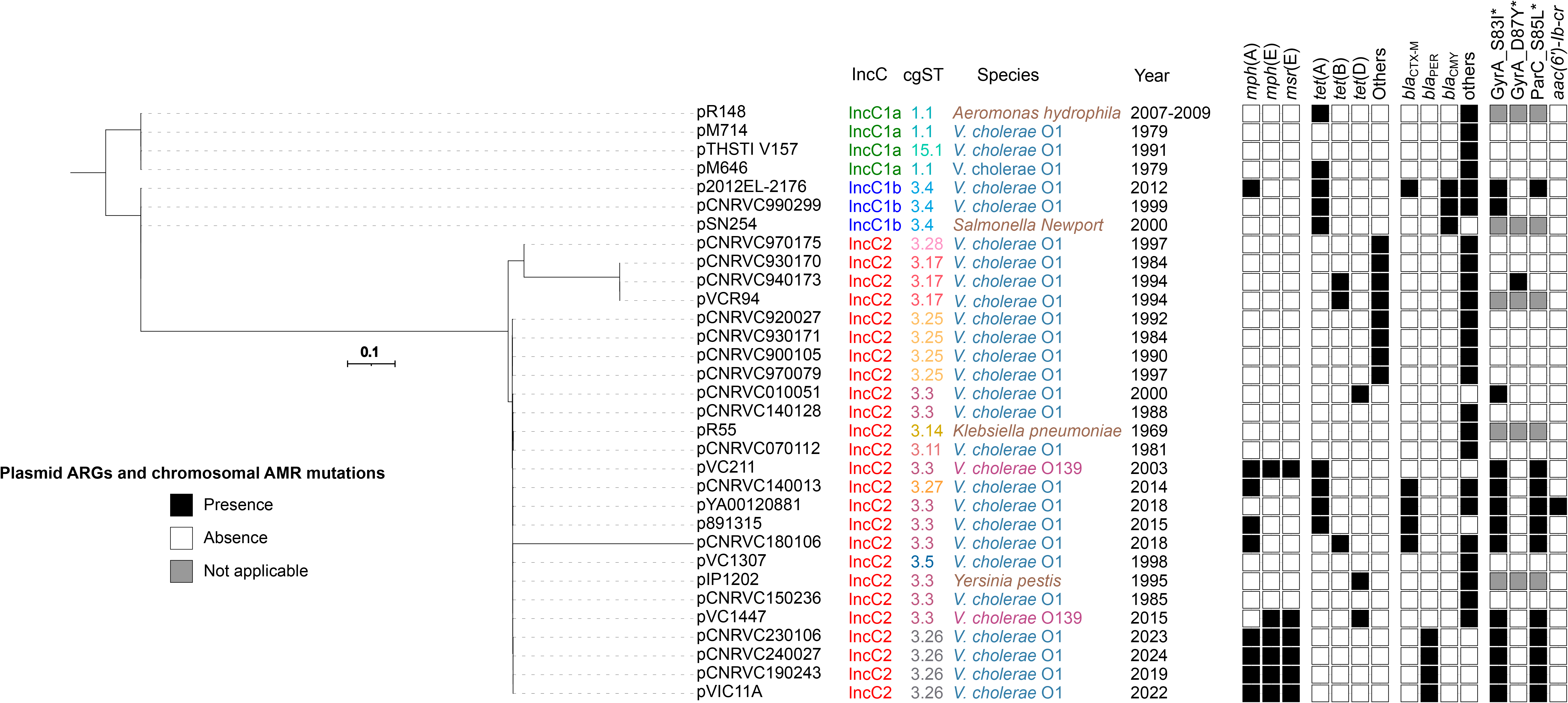
Maximum-likelihood phylogeny of 32 IncC plasmid backbones. Plasmid name, IncC type, cgST, bacterial host species, and year of isolation are indicated. The scale bar indicates the number of nucleotide substitutions per variable site. The matrix shows, from left to right, the presence or absence of plasmid-borne antimicrobial resistance (AMR) genes (ARGs) associated with azithromycin resistance (*mph*(A), *mph*(E), and *msr*(E)), tetracycline resistance (*tet*(A), *tet*(B), *tet*(D), and other *tet* genes), β-lactam resistance (*bla*_CTX-M_, *bla*_PER_, *bla*_CMY_, and other *bla* genes), and fluoroquinolone resistance (*aac(6’)-Ib-cr*). Asterisks indicate chromosomal mutations associated with fluoroquinolone resistance in *V. cholerae* strains harboring the corresponding plasmids. The exact ARG composition is provided in Supplementary Data 2.

In the global phylogenetic context, all IncC plasmids identified in African 7PET isolates were IncC2. They were mainly found in 7PET sublineages in which IncC plasmids were frequently observed, including AFR1, AFR5, AFR6, and AFR13, and less frequently in AFR7, AFR10, and AFR11 (Fig. 1). In contrast, IncC plasmids from Asian 7PET isolates included both IncC1a and IncC2 types in waves 1 and 2, whereas wave 3 isolates carried IncC1b or IncC2 plasmids. Notably, the three IncC2 plasmids from wave 2 isolates collected in China included two from *V. cholerae* O139 (Supplementary Data 2).

### High diversity of ARGs in the A/C plasmids studied

The 32 IncC plasmids studied and pRA1 (IncA) carried 62 distinct ARGs, conferring resistance to aminoglycosides, chloramphenicol, macrolides, rifampicin, β-lactams, sulfonamides, trimethoprim, and tetracyclines, with 2–16 ARGs per plasmid (mean and median, 8). *sul1* was the most prevalent ARG, detected in 27 plasmids (81.8%), followed by *sul2* in 25 (75.8%) (Supplementary Data 9 and 10). Among the 7PET *V. cholerae* plasmids, *sul2* was detected in IncC2 and IncC1b but not in IncC1a plasmids (Supplementary Data 10).

Among the 28 IncC plasmids from 7PET *V. cholerae*, 18 (64.3%) carried tetracycline resistance determinants. Tetracyclines are first-line antibiotics for the treatment of cholera, making the widespread occurrence of tetracycline resistance determinants of particular clinical relevance. However, the presence of these resistance determinants did not consistently correspond to phenotypic resistance: isolates carrying only *tet*(C) or *tet*(G) remained susceptible to tetracyclines (MICs, 1–4 mg/L), whereas those carrying *tet*(A), *tet*(B), or *tet*(D) were resistant or intermediately resistant (MICs, 8–32 mg/L) (Supplementary Data 2). These observations suggest that the phenotypic expression of tetracycline resistance varies according to the determinant carried and that the presence of an ARG alone may not reliably predict the resistance phenotype.

Among the 28 IncC plasmids from 7PET *V. cholerae*, 10 (35.7%) carried macrolide resistance determinants. Macrolides, including azithromycin, are used as second-line antibiotics for cholera treatment. Six plasmids (21.4%) carried resistance determinants for both macrolides and tetracyclines. Only isolate CNRVC230106 (Kenya, 2023), carrying *mph*(A), *mph*(E), and *msr*(E) on pCNRVC230106, showed phenotypic resistance to azithromycin (MIC >64 mg/L). In contrast, isolates 89315 (Zimbabwe, 2015) and CNRVC140013 (Democratic Republic of the Congo, 2014), each carrying a single copy of *mph*(A), and isolate CNRVC180106 (Somalia, 2018), carrying two copies of *mph*(A), remained susceptible to azithromycin (MIC, 1 mg/L).

Resistance to ciprofloxacin, another second-line antibiotic used for cholera treatment, is usually associated with mutations in the quinolone-resistance-determining regions (QRDRs) of the chromosomally encoded GyrA and ParC proteins, particularly the GyrA S83I and ParC S85L substitutions (Supplementary Data 2). This combination of mutations was previously reported in all isolates belonging to the AFR11–AFR15 sublineages ^7^. In our dataset, among the 27 7PET isolates for which chromosome sequences were available, 11 carried both substitutions. In contrast, plasmid-mediated quinolone resistance was rare, with *aac*(6’)-*Ib-cr* detected in only one plasmid, pYA00120881, carried by a strain associated with a cholera outbreak in Zimbabwe in 2018.

β-lactam antibiotics are not considered first-line treatments for cholera, but β-lactamase genes were widespread, with 17 distinct genes identified in 26 of the 28 IncC plasmids from 7PET *V. cholerae*. Ten plasmids (35.7%) carried β-lactamase genes associated with resistance to both ampicillin and third-generation cephalosporins, including ESBLs (*bla*SHV-12 *, bla*_CTX-M-2_, *bla*_CTX-M-15_, *bla*_CTX-M-202_, *bla*_PER-7_) and cephamycinase genes (*bla*_CMY-2_ and *bla*_CMY_). Notably, *bla*_CMY-2_ and *bla*_CMY_ (encoding a CMY-2-like β-lactamase with the Y219F amino-acid substitution) were detected exclusively in IncC1b plasmids and were absent from IncC2 and IncC1a plasmids. One plasmid, pCNRVC970079, from a strain isolated in the Central African Republic in 1997, carried *bla*_TEM_ (encoding a TEM-1-like β-lactamase with an L19F substitution) and a *bla*_SHV_-family gene (annotated as *bla*_SHV-12_ by ResFinder and *bla*_SHV-2a_ by AMRFinderPlus, Supplementary Data 4), but the corresponding isolate did not display phenotypic resistance to third-generation cephalosporins.

### Site-specific integration of AMR islands within a conserved IncC backbone

The IncC1 and IncC2 backbones are approximately 127.8 kb and 129.2 kb long, respectively, as deduced from analyses of pR148 and pR55 after removal of their AMR-associated regions (ARI-A from pR148 and Tn*6187* and GI*sul2* from pR55) ^16,57^. Notably, Tn*6187* is an unusual composite transposon structure containing two mercury resistance operons, two transposition modules related to Tn*21* and Tn*1696*, and an In0-type integron ^57^. Using the IncC2 pR55 as a reference, comparison with the other IncC plasmids in our dataset identified 28 distinct variable regions where genetic modules were present in one or more plasmids, either inserted into or replacing portions of the backbone, sometimes with additional modification of the surrounding sequence (Supplementary Data 11). These variable regions included AMR-associated modules (11 regions), ISs or transposons lacking ARGs (12 regions), and cryptic regions (five regions) containing, for example, hypothetical protein-coding sequences or CRISPR variants (Supplementary Data 11, Figs. 4 and 5). IncC2 plasmids contained more variable regions associated with ARG-bearing genetic structures (nine regions) than IncC1a (one region) or IncC1b (three regions) plasmids. Thus, although the ARG repertoire of *V. cholerae* IncC plasmids was diverse, resistance-associated genetic structures were frequently found at specific positions within the conserved IncC backbone.

### AMR islands show high plasticity and intricate evolutionary backgrounds

The principal AMR islands identified in our dataset were ARI-B and ARI-A. ARI-B was present in all 25 IncC2 plasmids (Supplementary Fig. 1) and three IncC1b plasmids (Figs. 4 and 5), whereas ARI-A was identified in IncC1a and IncC1b plasmids (Figs. 4 and 6).

**Figure 4:**
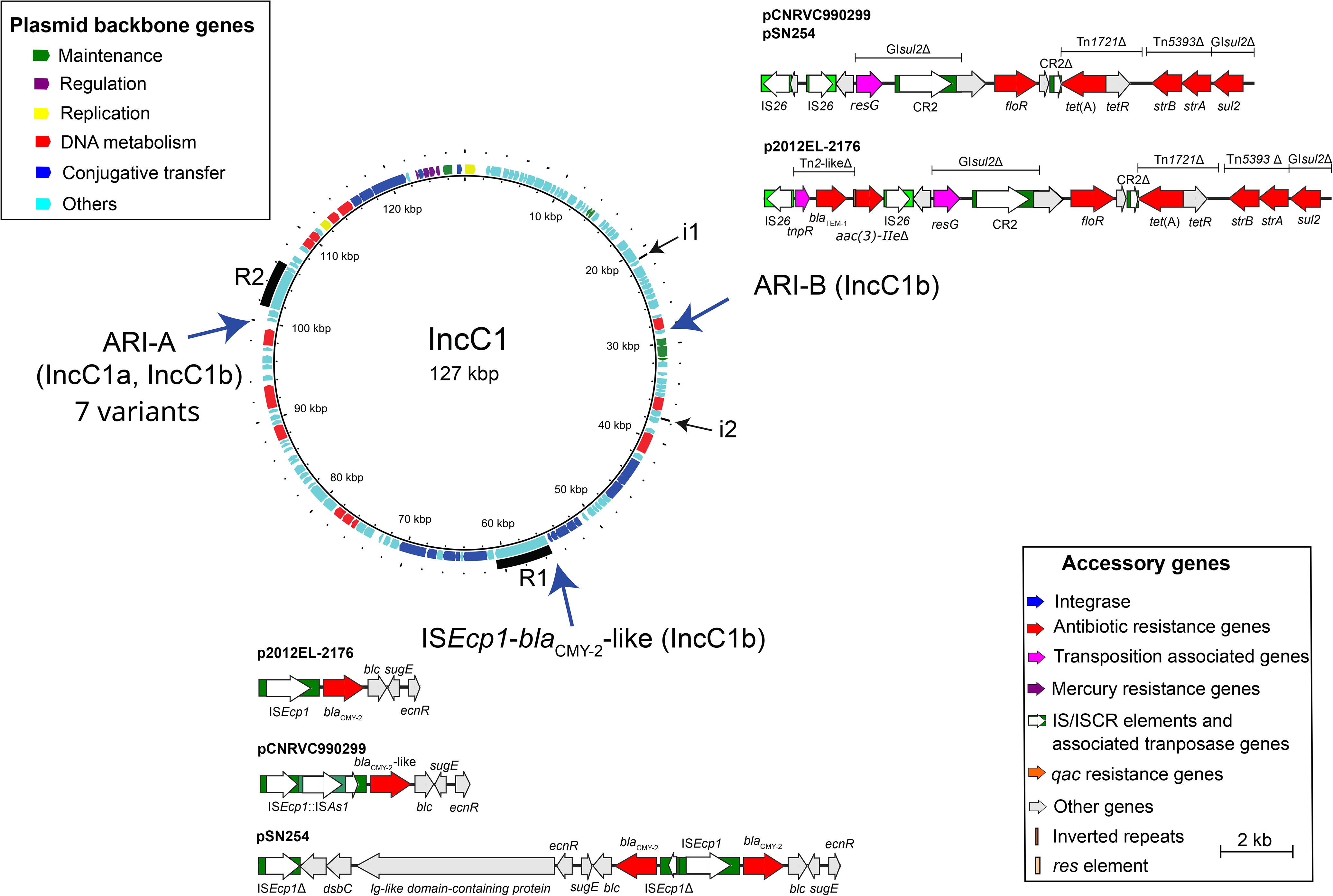
Insertions in IncC1 plasmids. Circles, from innermost to outermost, indicate: (1) the nucleotide positions of the IncC1 plasmid backbone derived from pR148 (GenBank accession no. JX141473.1); (2) backbone genes annotated as shown in the “Plasmid backbone genes” key. The displayed genes include those encoding proteins of more than 100 amino acids, as well as selected genes encoding shorter proteins with known functions; and (3) the locations of the i1, i2, R1, and R2 segments used to classify IncC plasmids as IncC1 or IncC2. The arrows indicate the sites of integration of antibiotic resistance structures in IncC1a and IncC1b plasmids. Different antibiotic resistance structures and their accessory genes are shown adjacent to the arrows and colored according to the “Accessory genes” key. Class 1 integrons were identified based on their cassette arrays. ARI-A islands are shown in Figure 6. The scale bar indicates the size of accessory structures.

**Figure 5:**
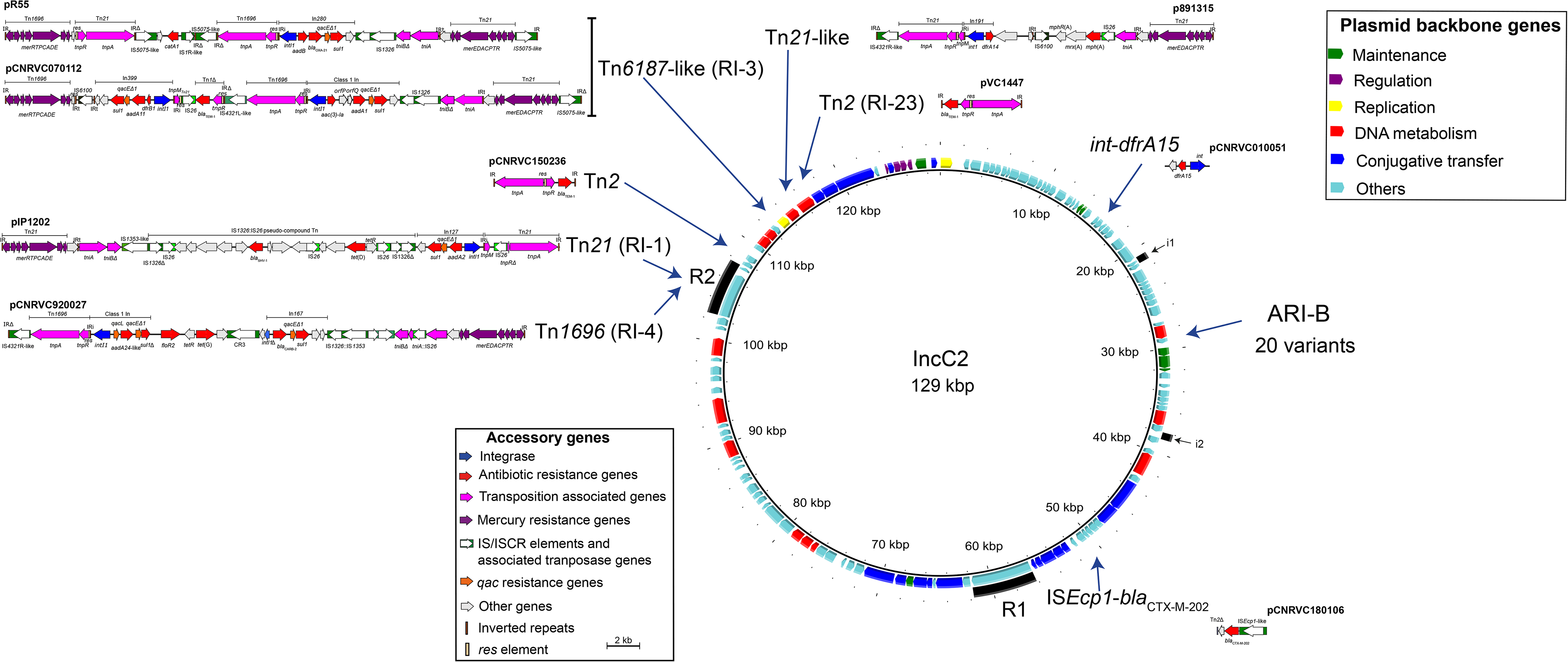
Insertions in IncC2 plasmids. Circles, from innermost to outermost, indicate: (1) the nucleotide positions of the IncC2 plasmid backbone derived from pR55 (GenBank accession no. JQ010984.1); (2) backbone genes annotated as shown in the “Plasmid backbone genes” key. The displayed genes include those encoding proteins of more than 100 amino acids, as well as selected genes encoding shorter proteins with known functions; and (3) the locations of the i1, i2, R1, and R2 segments used to classify IncC plasmids as IncC1 or IncC2. The arrows indicate the sites of integration of resistance islands in IncC2 plasmids. Different antibiotic resistance structures and their accessory genes are shown adjacent to the arrows and colored according to the “Accessory genes” key. Class 1 integrons were identified based on their cassette arrays. IRi and IRt are indicated where identified. RI refers to resistance islands as defined by Ambrose et al. ^16^. ARI-B islands are shown in Supplementary Fig. 1. The scale bar indicates the size of accessory structures.

**Figure 6:**
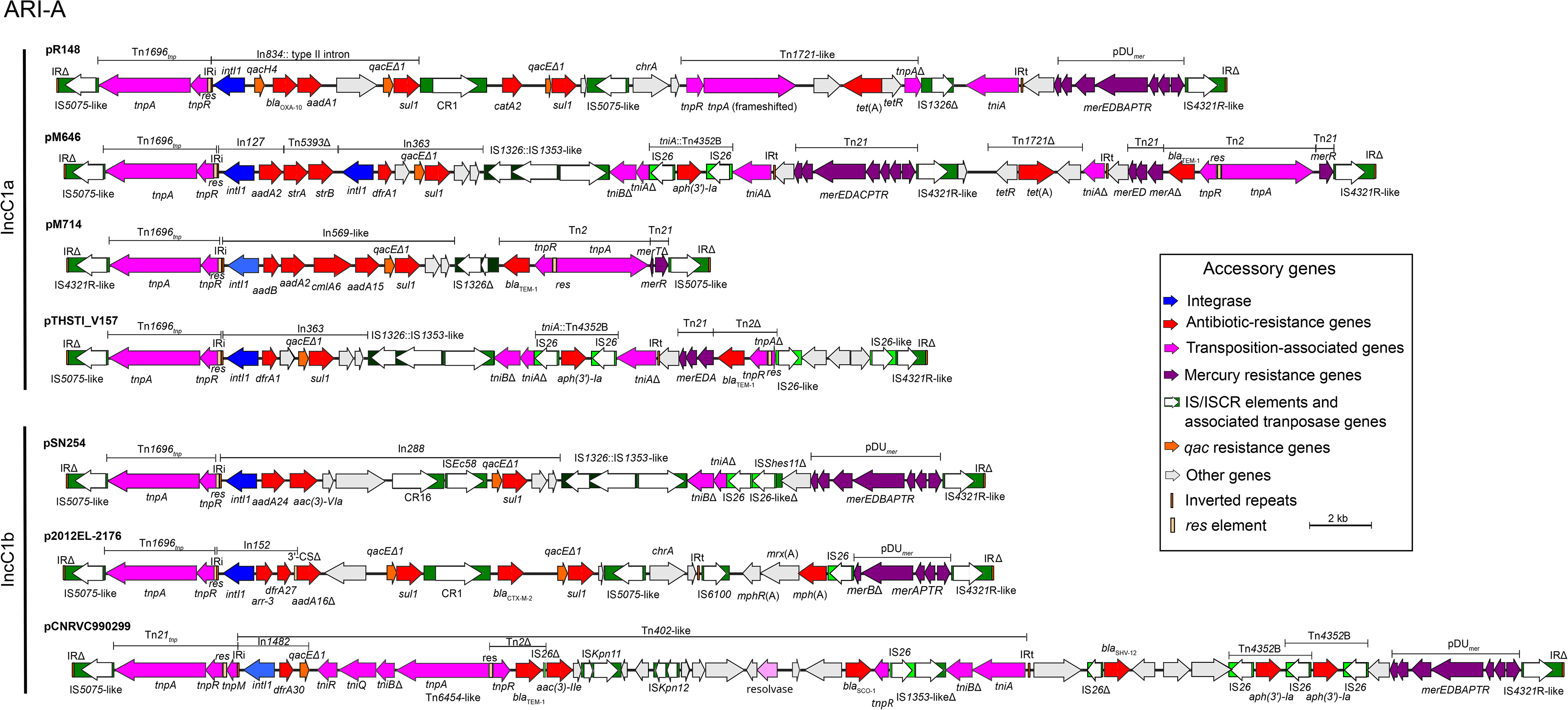
ARI-A structures in IncC1 plasmids. Arrows indicate gene length and orientation, with gene names shown above or below the arrows and colors corresponding to the associated key. The boundaries of retrieved modules are shown above the genomic region. Boxes in different shades of green represent insertion sequences (ISs) and common regions (CRs), with white arrows indicating the direction of transposition-associated genes. Class 1 integrons were identified based on their cassette arrays. IRi and IRt are indicated where identified. The scale bar indicates the size of accessory structures.

ARI-B showed extensive structural diversity while retaining features derived from the 15,460-bp genomic island GI*sul2* (GenBank accession no. KX709966.1, coordinates 29,940–45,399), including *sul2*. Its size ranged from 7.3 kb in pCNRVC010051 to 54.5 kb in pCNRVC140128, and individual ARI-B islands contained up to 16 ARGs interspersed with ISs, transposon remnants, transposons, and integrons (Supplementary Fig. 1). Several resistance determinants were embedded within recognizable mobile structures, including *strA* and *strB* within a partial Tn*5393*, *tet*(A) and *tet*(R) within a partial Tn*1721*, and *bla*_TEM_ variants within Tn*2*-like structures. *bla*_CTX-M-15_ was usually associated with IS*Ecp1*, while macrolide resistance genes were found within an IS*6100*-associated region containing *mphR*(A), *mrx*(A), and *mph*(A). Several structures within ARI-B were recurrent across multiple plasmids and 7PET sublineages. For example, ARG-containing structures comprising eight genes (*strA*, *strB*, *bla*_CARB-4_, *catA1*, *sul1*, *sul2*, *tet*(C), *dfrA15*) were identified in six IncC2 plasmids belonging to different 7PET sublineages. These structures included In*192*, an IS*26*-flanked pseudocompound transposon carrying *tet*(C), IS*1*R-*catA1*-Tn*21*_tnp_, and another putative IS*26*-flanked pseudocompound transposon containing two truncated Tn*5393*-associated *tnpR* sequences flanking a region comprising *tnsA*, *tnsB*, *bla*_CARB-4_, *strB*, *strA*, and an IS*Alw27*-like element. In some plasmids, this IS*Alw27*-like element was interrupted by IS*Pmi2*. These six plasmids were identified in AFR5 isolates (pCNRVC930170, pCNRVC940173, and pVCR94) ^26^, AFR1 isolates (pCNRVC900105 and pCNRVC930171), and an AFR6 isolate (pCNRVC970175). Although these plasmids showed minor rearrangements and remodeling in the ARI-B region, they maintained overall conservation of this resistance structure. These plasmids were recovered between 1984 and 1997 and belonged to three different cgSTs (3.17, 3.28, and 3.25), with backbone SNVs ranging from 0 to 962, indicating substantial divergence among some of the plasmids. Despite this backbone diversity, they shared closely related *tet*(C)-carrying structures, suggesting that different IncC plasmids may have independently acquired this structure on multiple occasions, followed by local structural remodeling. Historically, the emergence of tetracycline-resistant *V. cholerae* during the 1977–1978 cholera outbreak in Tanzania was associated with extensive prophylactic use of tetracycline (1.79 metric tons used over five months) and the emergence of an IncC plasmid in outbreak strain ^59^. This episode provides historical context for the early selection and dissemination of tetracycline resistance in 7PET. Interestingly, pCNRVC940173 and pVCR94, both recovered in 1994, carried a second tetracycline resistance gene, *tet*(B), in addition to *tet*(C). The *tet*(B) gene was also identified in pCNRVC180106, recovered in 2018. In all three plasmids, *tet*(B) was located within truncated Tn*10* derivatives, which were additionally interrupted by an IS*Kpn26*-like element in pCNRVC940173 and pVCR94. Despite these structural differences, the *tet*(B)-containing regions shared homologous flanking sequences, including genes predicted to be involved in membrane transport, metal homeostasis, and redox processes upstream, and a *catA1*–Tn*21*-associated *tnp* region downstream (Supplementary Fig. 1).

Class 1 integrons were also frequent within ARI-B. The In*192* integron (GQ280261), carrying *dfrA15*, was identified in seven ARI-B islands, whereas pCNRVC140128 carried a class 2 integron. Transposition modules related to Tn*21* and Tn*1696*, without their associated mercury-resistance modules, were present in many ARI-B islands. An exception was pCNRVC140128, which carried a hybrid structure combining a Tn*21* transposition module with a mercury-resistance module derived from Tn*1696*. Among the 28 ARI-B islands, 20 started with IS*26*, whereas the remaining eight started with IS*1326*, IS*6100*, CR1, the integrase associated with GI*sul2*, or a remnant type IV secretion system gene cluster (*virB8–virB11*) (Fig. 4 and Supplementary Fig. 1).

Despite this extensive diversity, *sul2* was the only ARG detected exclusively within ARI-B, occurring in 24 of the 28 ARI-B islands (Fig. 4, Supplementary Fig. 1, and Supplementary Data 12). ARI-B was integrated at the same backbone position in both IncC2 and IncC1b plasmids. None of the plasmids contained an intact copy of GI*sul2*; however, most retained its 3′ region containing *sul2* and the mobile element CR2 ^60^. A structure containing both ends of GI*sul2*, with the central region replaced by a fragment containing the phenicol-resistance gene *floR*, was observed in only two plasmids, pR55 and pCNRVC070112. These plasmids lacked detectable deletions in the surrounding backbone (Fig. 2). In contrast, ARI-B in several other plasmids was associated with deletions of adjacent backbone sequences, including the i1 region, providing a likely explanation for the observed variation in this segment (Fig. 2 and Supplementary Figs. 2 and 3).

Substantial differences in ARI-B content were observed among 20 IncC2 (Supplementary Fig. 1) and two IncC1b plasmids (Fig. 4), indicating extensive remodeling of this region. Nevertheless, closely related ARI-B architectures were also observed across different bacterial species and epidemiological contexts. The ARI-B islands of the IncC2 plasmids pR55, from a *K. pneumoniae* isolate collected in France in 1969, and pCNRVC070112, from a 7PET AFR1 *V. cholerae* O1 isolate collected in Algeria in 1981, were identical except for three apparent indels within homopolymeric regions, likely reflecting sequencing or assembly errors in the previously published pR55 sequence. Similarly, the ARI-B islands of the IncC1b plasmids pSN254, from *Salmonella* collected in the USA in 2000, and pCNRVC990299, from a wave 3 7PET *V. cholerae* O1 isolate collected in Afghanistan in 1999, were similar (99% identity). More broadly, pR55-like and pSN254-like plasmids, defined by >95% query coverage and >99% sequence identity in BLAST, are found in multiple bacterial species and are frequently represented in sequence databases.

Additional evidence for the conservation and dissemination of related AMR structures was provided by the similar ARI-B architectures observed in pCNRVC150236, from a 1985 AFR5 *V. cholerae* O1 isolate in Somalia, pIP1202, from *Y. pestis* collected in Madagascar in 1995 ^55^, and pVC1447, from *V. cholerae* O139 collected in China in 2015 ^61^. These regions shared a common overall organization, with type IV secretion system-associated genes followed by potential pseudocompound transposons carrying different resistance determinants and a GI*sul2*-derived structure at the distal end (Supplementary Fig. 1). Notably, pIP1202 and pCNRVC010051, both IncC2 cgST3.3 plasmids, were recovered in Madagascar in 1995 from *Y. pestis*, the causative agent of plague, and in 2000 from *V. cholerae* O1, respectively. Despite their closely related plasmid backbones, they carried distinct ARG repertoires, illustrating the capacity of a shared plasmid lineage to circulate among major human pathogens while acquiring and remodeling different resistance cargoes over time.

Within 7PET *V. cholerae*, ARI-B structures were also sometimes conserved among plasmids from the same sublineage, as observed in AFR13 (pVIC11-A, pCNRVC230106, pCNRVC240027, and pCNRVC190243), whereas other plasmids shared only individual sequence blocks, as observed for pCNRVC930170 and pCNRVC940173 (Supplementary Fig. 1). Together, these observations indicate that ARI-B combines a conserved genomic framework with extensive and recurrent remodeling of its resistance cargo, allowing related IncC plasmids to maintain shared structural features while acquiring different combinations of AMR genes across hosts, lineages, and time.

ARI-A ranged from 20.9 kb in pM714 to 48.5 kb in pCNRVC990299 and carried three to eight ARGs. All seven ARI-A regions were bounded by terminal inverted repeats interrupted by either IS*5075* or IS*4321*R (Fig. 6), potentially limiting their further transpositional mobility ^17^. ARI-A consisted of complex derivatives of Tn*1696*, with Tn*1696*-related transposition modules in six plasmids and a Tn*21*-related module in pCNRVC990299 ^62^. The mercury-resistance module was derived from pDU*mer,* similar to that described in plasmid pRMH760 (NC_023898.1:140518-144785) in the three IncC1b and one IncC1a plasmids, and from Tn*21* in the three *V. cholerae* IncC1a plasmids. The Tn*21*-derived mercury modules were incomplete except in pM646, which contained one complete and one incomplete copy, potentially resulting from recombination between the two IS*4321*R elements.

ARI-A also contained a mosaic of class 1 integrons and transposons. Most class 1 integrons, except those of pM714 and p2012EL-2176, were associated with truncated *tni* regions derived from Tn*402*. Embedded transposons included Tn*2*, carrying *bla*_TEM-1_, in four ARI-A regions; Tn*1721*, carrying *tet*(A), in two; and the IS*26*-flanked pseudocompound transposon Tn*4352*B, carrying *aph(3’)-Ia*, in three. Notably, pCNRVC990299 contained a tandem array of two overlapping Tn*4352*B units in which three IS*26* copies were aligned in the same orientation, consistent with sharing of the central IS*26* element between the two units. This structure could have arisen through IS*26*-mediated targeted conservative cointegration, whereby an existing IS*26* copy serves as a target for the integration of an additional IS*26*-associated segment ^63^. None of the seven ARI-A regions showed an identical overall structure, highlighting their substantial structural diversity and supporting the high structural plasticity and rapid evolution previously reported for ARI-A ^16^.

While some insertions (like ARI-B and ARI-A) likely occur within specific regions of the IncC backbone, we also identified other less common antibiotic resistance structures. pCNRVC070112, like pR55, contains a resistance island (RI-3 according to Ambrose et al. ^16^), inserted between *yacC* and *ter* genes of the IncC2 backbone (Fig. 5). This island shares similarities with Tn*6187* from pR55 ^57^, with two mercury resistance modules related to Tn*21* and Tn*1696*. In contrast to Tn*6187* in pR55, it has a single transposition module related to Tn*1696* and lacks the Tn*21* module; it also contains two class 1 integrons instead of one in pR55, with distinct cassette arrays, and a deleted Tn*1* with the *bla*_TEM-1_ gene. pCNRVC920027 contains a resistance island (RI-4 according to Ambrose et al. ^16^) inserted into the *rhs2* gene of the IncC2 backbone (Fig. 5). It consists of a Tn*1696*-related transposon with a Tn*1696* transposition module and a distinct mercury resistance module. Unlike canonical Tn*1696* (U12338.3), this transposon carries two class 1 integrons, one of which harbors a new cassette array (*qacL* and a new *aadA* variant) and the other one, *bla*_CARB-2_. Interestingly, CR3 is located between these two integrons and is associated with *tet*(G) and the *floR* variant (*floR2*). CR3, along with *tet*(G) and *floR*, was generally identified within SGI1 and was also located between two class 1 integrons (one with *aadA2* and the other with *bla*_CARB-2_) ^64^. Comparing SGI1 (AF261825.2) with pCNRVC920027, the 12,565-bp region (pCNRVC920027 coordinates, 119265–131820), spanning from *aadA* in the first integron to the end of the second integron, showed 98% identity, highlighting similarities between resistance modules of SGI1 and the pCNRVC920027 RI-4. While pIP1202 has Tn*21* with an *aadA2* cassette array inserted into the IncC2 backbone within the *rhs2* gene (defined as RI-1 by Ambrose et al. ^16^), p891315 has another Tn*21*-related transposon inserted into a different IncC2 backbone gene (*uvrD*) (Fig. 5). This p891315 has a different cassette array (*dfrA14*) from canonical Tn*21* (AF071413.3) and lacks both a 3′-conserved region and common Tn*21* IS elements, such as IS*1326*. Unlike canonical Tn*21*, an IS*4321*R-like element is also inserted into one of the IRs of Tn*21* p891315. Within this p891315 Tn*21*-like element is an *mph*(A) region flanked by two ISs: IS*6100*, *mphR*(A), *mrx*(A), *mph*(A), and IS*26*.

Small antibiotic resistance structures were also found scattered in the IncC2 backbone, such as Tn*2* inserted in two different positions in the IncC2 backbone, IS*Ecp1*-*bla*_CTX-M-202_ and *int*-*dfrA15* (Fig. 5). The *bla*_CMY-2_ gene (and its variants), an *ampC*-type β-lactamase gene conferring resistance to third-generation cephalosporins, was detected exclusively in IncC1b plasmids, within IS*Ecp1*-associated transposition units (TUs), and was absent from the IncC2 and IncC1a plasmids (Fig. 4), consistent with previous observations ^16^. IS*Ecp1* is known to mobilize DNA adjacent to its right-hand end as a TU ^65^, and the three IncC1b plasmids carrying *bla*_CMY_ variants showed related but distinct TU structures. The IS*Ecp1*-*bla*_CMY-2_ TU in p2012EL-2176 was essentially identical to the corresponding TU in pAR060302, an IncC plasmid recovered from *E. coli* isolated from a dairy cow in 2002 ^66^. In pCNRVC990299, the TU contained an IS*Ecp1* element interrupted by IS*As1* and an unnumbered *bla*_CMY_ variant that was 99% identical to *bla*_CMY-2_, suggesting subsequent modification of the original TU. The pSN254 plasmid contained two copies of *bla*_CMY-2_ and duplicated regions within a more complex IS*Ecp1*-associated structure. This structure shares a segment with Tn*7241* (KY437729.1), a complex transposon formed by rearrangement of an IS*Ecp1*-*bla*_CMY-6_ TU and its capture between two IS*10* elements ^67^, suggesting that pSN254 may represent an ancestral or related structure in the evolutionary history of Tn*7241*. Together, these observations illustrate the structural diversification of IS*Ecp1*-associated *bla*_CMY_ TUs within IncC1b plasmids.

Across the 32 IncC plasmids, we identified 20 class 1 and one class 2 integrons based on their cassette arrays, including two class 1 integrons with new cassette arrays not previously described (Supplementary Data 13).

### A potential CRISPR locus within a *V. cholerae* IncC plasmid

CRISPR arrays are predominantly chromosome-encoded, whereas their occurrence on plasmids is comparatively uncommon and heterogeneous ^68,69^. A 1.7-kb putative CRISPR array (GenBank accession no. PZ168218, coordinates 148,632–150,309) was identified using CRISPRCasFinder in pCNRVC180106, an IncC2 plasmid obtained from a 7PET AFR13 *V. cholerae* O1 isolate collected in Somalia in 2018. This array comprised 28 direct repeats (DRs) and 27 spacers. The array was located immediately upstream of *csf4*, which encodes a CRISPR-associated DEAD/DEAH-box helicase (coordinates 150,562–152,259). BLAST analysis of these spacers against the 1,530 7PET genomes in our phylogenetic dataset identified five spacers (4, 5, 6, 19, and 23) that showed full-length matches to sequences present specifically in 75 wave 2 genomes carrying GI-15 ^7^, with one to four nucleotide differences (Supplementary Data 14 and 15). Three additional spacers (9, 21, and 22) partially matched GI-15. Six spacers (4, 5, 6, 10, 19, and 23) also showed full-length matches to sequences within genes of *Salmonella* Genomic Island 1 (SGI1) (GenBank accession nos. AF261825.2 and CP014358.1), including genes involved in SGI1 transfer, with zero to four nucleotide differences, whereas two additional spacers (9 and 22) partially matched SGI1 (AF261825.2, and CP014358.1) (Supplementary Data 14). SGI1 is a 42.4-kb integrative and mobilizable element carrying multiple ARGs, including *bla*_CARB-2_, *floR*, *aadA2*, *sul1*, and *tet*(G) ^70^. GI-15 is a related 28,172-bp genomic element that carries *aadA1* and *sul1* ^71^. Notably, none of the spacer-targeted sequences overlapped these ARGs in either SGI1 or GI-15. SGI1 and related elements can be mobilized in trans by IncC plasmids through the interaction of SgaCD encoded by SGI1 and AcaCD encoded by IncC ^72^. Conversely, several studies have reported antagonistic interactions between SGI1-like elements and A/C plasmids, including inhibition of plasmid transfer and, potentially, plasmid exclusion ^73–76^. Consistent with this, our analysis of 1,530 7PET genomes revealed an apparent mutual exclusivity between IncC plasmids and GI-15, with isolates generally carrying either an IncC plasmid or GI-15 but not both _7_. The presence of multiple spacers with matches to SGI1 and GI-15 within pCNRVC180106 is therefore notable in the context of this apparent antagonism. However, the putative CRISPR locus does not appear to be associated with a complete Cas region, raising the possibility that it is incomplete or inactive. Although the spacer content may reflect past interactions with SGI1-like elements, whether this locus represents a functional CRISPR system and whether it affects IncC plasmid stability or transfer in the presence of GI-15 remain to be determined.

### Most of the IncC plasmids tested are conjugative

Nine of the 10 IncC plasmids from 7PET *V. cholerae* donor strains tested were successfully transferred to *E. coli* K-12 J5 (Supplementary Data 6). pCNRVC970175 was not transferred, potentially owing to a ∼20-kb deletion in its backbone encompassing *ssb* and a *bet/exo* recombination module (Supplementary Fig. 4). The *bet* and *exo* genes encode recombination functions involved in DNA repair ^77^. The deletion of this region could potentially contribute to the failure of plasmid transfer or maintenance in the recipient, although the underlying mechanism remains uncertain.

The conjugation experiments also revealed differences between plasmid-borne resistance determinants and the observed phenotype in *V. cholerae*. The *V. cholerae* isolate carrying pCNRVC180106, which contains two copies of *mph*(A), was susceptible to azithromycin (MIC, 1 mg/L), whereas its *E. coli* transconjugants were resistant (MIC, 64 mg/L) (Supplementary Data 16). Similarly, the *V. cholerae* isolate carrying pCNRVC140013, which contains a single copy of *mph*(A), was susceptible to azithromycin (MIC, 1 mg/L), whereas its *E. coli* transconjugants were resistant (MICs, 8–32 mg/L). The *V. cholerae* isolate carrying p891315, which contains a single copy of *mph*(A), was also susceptible to azithromycin (MIC, 1 mg/L); no conjugation experiment was performed with this isolate. These observations suggest that expression of plasmid-borne macrolide resistance determinants may be influenced by the bacterial host background. However, the mechanisms underlying these differences remain unclear. They could involve differences in gene expression or regulation between *V. cholerae* and *E. coli*, although changes in plasmid structure or gene content during conjugation cannot be excluded because the transconjugants were not characterized by long-read sequencing.

The highly drug-resistant AFR13 *V. cholerae* isolates carrying cgST3.26 IncC2 plasmids with the three macrolide resistance determinants *mph*(A), *mph*(E), and *msr*(E) showed an azithromycin-resistant phenotype (MICs, 32–>64 mg/L). In contrast, the three *V. cholerae* isolates mentioned above, carrying only one or two copies of *mph*(A) as the only genomic determinant associated with azithromycin resistance, were nevertheless susceptible to azithromycin. Because these isolates also carried genomic determinants associated with resistance to tetracycline (plasmid-borne) and ciprofloxacin (chromosomally encoded), their genomic AMR profiles would have suggested resistance to the three oral antibiotics recommended for cholera treatment, whereas the observed azithromycin phenotype remained susceptible. This highlights the limitations of inferring antimicrobial susceptibility from ARG content alone and warrants further investigation of the host and genetic factors underlying the expression of plasmid-borne resistance determinants.

### AFR sublineages have acquired multiple IncC plasmid types on different occasions

Several 7PET sublineages, including AFR1, AFR5, and AFR13, acquired distinct IncC plasmids on multiple occasions (Fig. 1, Table 2). Among these, AFR13 provides a notable example of the subsequent geographic dissemination of an IncC2 plasmid-associated clone. An IncC2 plasmid carrying a highly drug-resistant profile emerged during the 2018–2019 cholera outbreak in Yemen and was subsequently detected across multiple geographic locations. Before this period, AFR13 isolates from Yemen had a relatively narrow resistance profile. By late 2018, an AFR13 clone carrying pCNRVC190243, which contained *bla*_PER-7_ and a macrolide-resistance region carrying *mph*(A), *mph*(E), and *msr*(E), emerged and subsequently spread to South Lebanon, Kenya, Tanzania, Comoros, and Mayotte ^12,30,32^. Closely related plasmids belonging to cgST3.26 differed by only 0–2 SNVs in their backbones (Supplementary Data 8), consistent with the dissemination of a highly conserved plasmid within this bacterial clone. The same clone has also been identified in travelers returning to England from Yemen, Kenya, and Zimbabwe ^78^, as well as in cholera cases in the United Kingdom and Germany associated with consumption of contaminated holy water imported from Ethiopia ^79^. The widespread dissemination of this 7PET clone carrying a highly drug-resistant IncC2 pCNRVC190243 represents a potential threat to cholera treatment, which could be further exacerbated if this clone were to acquire an additional tetracycline-resistance determinant, potentially compromising the effectiveness of currently recommended oral treatments. The AFR13 sublineage has also acquired other highly drug-resistant IncC plasmids in more geographically restricted settings. pYA00120881, carried by an epidemic strain from Zimbabwe in 2018, contained 16 ARGs, including *aac(6’)-Ib-cr*, the first reported plasmid-mediated quinolone-resistance determinant in *V. cholerae*, together with *tet*(A) ^31^. pCNRVC180106, recovered from a ciprofloxacin-resistant isolate from Somalia in 2018, carried 14 ARGs, including *tet*(B) and two copies of *mph*(A), but remained susceptible to azithromycin (see previous paragraph).

AFR5 provides a complementary example of repeated IncC acquisition over a longer period. This sublineage circulated in the African Great Lakes region during the 1980s and 1990s, and almost all sampled AFR5 isolates from 1984–1998 carried multidrug-resistant IncC plasmids ^7^. The closely related plasmids pCNRVC930170 (1984) and pCNRVC940173 (1994), both recovered in the Democratic Republic of the Congo 10 years apart, shared the same cgST (cgST3.17) and differed by only one SNV across their conserved backbones. They also shared three of four insertion sites for accessory genetic elements, including mobile genetic elements and resistance-associated structures. In contrast, their ARI-B regions had undergone multiple rearrangements, including insertions and deletions associated with IS elements, resulting in loss of several ARGs and acquisition of *tet*(B), while other resistance determinants were retained.

A closely related plasmid, pVCR94, was also identified in 1994, in neighboring Rwanda ^26^. In parallel, pCNRVC150236, recovered from another AFR5 isolate in Somalia in 1985, belonged to a different IncC2 backbone type (cgST3.3). Together, these observations indicate that AFR5 repeatedly acquired and maintained different IncC plasmids during its circulation across Central and Eastern Africa, while individual plasmids underwent substantial remodeling of their resistance cargoes.

In conclusion, IncC plasmids have contributed substantially to the acquisition, maintenance, and dissemination of antimicrobial resistance in 7PET *V. cholerae*. Their remarkably conserved backbones over almost five decades contrast with the extensive diversification of ARGs and associated genomic islands, demonstrating that a relatively stable plasmid scaffold can repeatedly acquire and remodel resistance cargo under changing selective pressures. The occurrence of closely related IncC plasmid backbones and resistance-associated structures in diverse bacterial species further illustrates their broad host range and the potential for interspecies dissemination of clinically important resistance determinants. The recent detection of a 142-kb A/C plasmid carrying *bla*_NDM-1_, *bla*_CMY-6_, and *bla*_DHA-7_ in six 7PET isolates from Gujarat, India, in 2019 ^80^, although its complete sequence was unavailable, underscores the continuing emergence of clinically important resistance within this plasmid family. These findings highlight the importance of One Health approaches, genomic surveillance, and antimicrobial stewardship to monitor and limit the dissemination of IncC plasmids and the resistance determinants they carry.

## AUTHOR CONTRIBUTIONS

FXW designed and oversaw the study. SIM, AMS, NB, CP, TR, MLQ, CR, and FXW selected and provided isolates or genomes with their basic metadata. EN and MG subcultured the bacteria, performed phenotypic experiments, and extracted DNA. RR, EN, MG, AMS, MLQ, CR, and FXW analyzed and/or interpreted the data. FD, MH, and FXW provided guidance to RR. RR wrote the manuscript, with a major contribution from FXW. All the authors contributed to the editing of the manuscript.

## ACKNOWLEDGEMENTS

This research was funded by the *Fondation Le Roch-Les Mousquetaires* (to F.-X.W); *Institut Pasteur* (to F.-X.W); *Santé publique France* (to F.-X.W); and by the French Government’s *Investissement d’Avenir* programme, *Laboratoire d’Excellence* ‘Integrative Biology of Emerging Infectious Diseases’ (grant no. ANR-10-LABX-62-IBEID to F.-X.W). R.R. was funded by a grant **“***bourses de Séjours Scientifiques de Haut Niveau (SSHN)*” from the French Embassy in Lebanon. The authors sincerely thank Alexandra Moura for her assistance in the annotation of integrons in this manuscript. They also thank the IncA/C cgPMLST team at PubMLST for assigning allele and sequence type numbers to the newly submitted sequences.

## USE OF ARTIFICIAL INTELLIGENCE TOOLS

ChatGPT (OpenAI) was used solely for English-language editing of the manuscript, including grammar, syntax, wording and readability. The authors reviewed and approved all edited text and remain fully responsible for the content, scientific accuracy and conclusions of the manuscript.

## REFERENCES

1. Kanungo, S., Azman, A. S., Ramamurthy, T., Deen, J. & Dutta, S. Cholera. Lancet 399, 1429–1440 (2022).

2. World Health Organization. Cholera. https://www.who.int/news-room/fact-sheets/detail/cholera (2026).

3. Vega Ocasio, D., et al. Cholera outbreak -Haiti, September 2022-January 2023. MMWR Morb. Mortal. Wkly. Rep. 72, 21–25 (2023).

4. World Health Organization. Cholera situation in Yemen, April 2021. https://reliefweb.int/report/yemen/cholera-situation-yemen-april-2021 (2022).

5. Sudanese American Physicians Association. Evaluation and lessons learned: cholera outbreak response in Khartoum and Tawila, Sudan. https://reliefweb.int/report/sudan/evaluation-and-lessons-learned-cholera-outbreak-response-khartoum-and-tawila-sudan (2025).

6. Rouard, C., Njamkepo, E., Quilici, M.-L. & Weill, F.-X. Contribution of microbial genomics to cholera epidemiology. C. R. Biol. 345, 37–56 (2022).

7. Weill, F.-X. et al. Genomic history of the seventh pandemic of cholera in Africa. Science 358, 785–789 (2017).

8. Mutreja, A. et al. Evidence for several waves of global transmission in the seventh cholera pandemic. Nature 477, 462–465 (2011).

9. Weill, F.-X. et al. Genomic insights into the 2016-2017 cholera epidemic in Yemen. Nature 565, 230–233 (2019).

10. Benamrouche, N. et al. Outbreak of Imported Seventh Pandemic *Vibrio cholerae* O1 El Tor, Algeria, 2018. Emerg. Infect. Dis. 28, 1241–1245 (2022).

11. Smith, A. M. et al. Imported Cholera Cases, South Africa, 2023. Emerg. Infect. Dis. 29, 1687–1690 (2023).

12. Lassalle, F. et al. Genomic epidemiology reveals multidrug resistant plasmid spread between *Vibrio cholerae* lineages in Yemen. Nat. Microbiol. 8, 1787–1798 (2023).

13. Saha, D. et al. Single-dose azithromycin for the treatment of cholera in adults. N. Engl. J. Med. 354, 2452–2462 (2006).

14. Jones, E. H. et al. *Vibrio* Infections and Surveillance in Maryland, 2002–2008. Public Health Rep. 128, 537–545 (2013).

15. Ambrose, S. J., Harmer, C. J. & Hall, R. M. Compatibility and entry exclusion of IncA and IncC plasmids revisited: IncA and IncC plasmids are compatible. Plasmid 96–97, 7–12 (2018).

16. Ambrose, S. J., Harmer, C. J. & Hall, R. M. Evolution and typing of IncC plasmids contributing to antibiotic resistance in Gram-negative bacteria. Plasmid 99, 40–55 (2018).

17. Harmer, C. J. & Hall, R. M. The A to Z of A/C plasmids. Plasmid 80, 63–82 (2015).

18. Maimone, F. et al. Clonal spread of multiply resistant strains of *Vibrio cholerae* O1 in Somalia. J. Infect. Dis. 153, 802–803 (1986).

19. Threlfall, E. J. & Rowe, B. *Vibrio cholerae* El Tor acquires plasmid-encoded resistance to gentamicin. Lancet 1, 42 (1982).

20. Hedges, R. W., Vialard, J. L., Pearson, N. J. & O’Grady, F. R plasmids from Asian strains of *Vibrio cholerae*. Antimicrob. Agents Chemother. 11, 585–588 (1977).

21. Tabtieng, R. et al. An epidemic of *Vibrio cholerae* El tor Inaba resistant to several antibiotics with a conjugative group C plasmid coding for type II dihydrofolate reductase in Thailand. Am. J. Trop. Med. Hyg. 41, 680–686 (1989).

22. Rahal, K., Gerbaud, G. & Bouanchaud, D. H. Stability of R plasmids belonging to different incompatibility groups in *Vibrio cholerae* ‘Eltor’. Ann. Microbiol. (Paris) 129, 409–414 (1978).

23. Hochhut, B., Beaber, J. W., Woodgate, R. & Waldor, M. K. Formation of chromosomal tandem arrays of the SXT element and R391, two conjugative chromosomally integrating elements that share an attachment site. J. Bacteriol. 183, 1124–1132 (2001).

24. Wozniak, R. A. F. et al. Comparative ICE genomics: insights into the evolution of the SXT/R391 family of ICEs. PLoS Genet. 5, e1000786 (2009).

25. Blokesch, M. & Seed, K. D. Lineage-specific defence systems of pandemic *Vibrio cholerae*. Philos. Trans. R. Soc. Lond. B Biol. Sci. 380, 20240076 (2025).

26. Carraro, N. et al. Development of pVCR94ΔX from *Vibrio cholerae*, a prototype for studying multidrug resistant IncA/C conjugative plasmids. Front. Microbiol. 5, 44 (2014).

27. Loeff, L. et al. Molecular mechanism of plasmid elimination by the DdmDE defense system. Science 385, 188–194 (2024).

28. Jaskólska, M., Adams, D. W. & Blokesch, M. Two defence systems eliminate plasmids from seventh pandemic *Vibrio cholerae*. Nature 604, 323–329 (2022).

29. Blokesch, M. Defence systems encoded by core genomic islands of seventh pandemic *Vibrio cholerae*. Philos. Trans. R. Soc. Lond. B Biol. Sci. 380, 20240083 (2025).

30. Abou Fayad, A., et al. An unusual two-strain cholera outbreak in Lebanon, 2022-2023: a genomic epidemiology study. Nat. Commun. 15, 6963 (2024).

31. Mashe, T. et al. Highly Resistant Cholera Outbreak Strain in Zimbabwe. N. Engl. J. Med. 383, 687–689 (2020).

32. Rouard, C. et al. Long-distance spread of a highly drug-resistant epidemic cholera strain. N. Engl. J. Med. 391, 2271–2273 (2024).

33. Carraro, N., Rivard, N., Ceccarelli, D., Colwell, R. R. & Burrus, V. IncA/C conjugative plasmids mobilize a new family of multidrug resistance islands in clinical *Vibrio cholerae* non-O1/non-O139 isolates from Haiti. mBio 7, e00509–16 (2016).

34. Croucher, N. J. et al. Rapid phylogenetic analysis of large samples of recombinant bacterial whole genome sequences using Gubbins. Nucleic Acids Res. 43, e15 (2015).

35. Stamatakis, A. RAxML-VI-HPC: maximum likelihood-based phylogenetic analyses with thousands of taxa and mixed models. Bioinformatics 22, 2688–2690 (2006).

36. Letunic, I. & Bork, P. Interactive Tree Of Life (iTOL) v4: recent updates and new developments. Nucleic Acids Res. 47, W256–W259 (2019).

37. The European Committee on Antimicrobial Susceptibility Testing. Breakpoint tables for interpretation of MICs and zone diameters. version 16.0. http://www.eucast.org (2026).

38. Clinical and Laboratory Standards Institute. Methods for antimicrobial dilution and disk susceptibility testing of infrequently isolated or fastidious bacteria; approved guideline. 3rd Edition (M45Ed3E). (2010).

39. Clinical and Laboratory Standards Institute. Performance standards for antimicrobial susceptibility testing. 27th Ed. CLSI Supplement M100. (2017).

40. Wick, R. R., Judd, L. M., Gorrie, C. L. & Holt, K. E. Unicycler: resolving bacterial genome assemblies from short and long sequencing reads. PLoS Comput. Biol. 13, e1005595 (2017).

41. Carattoli, A., et al. *In silico* detection and typing of plasmids using PlasmidFinder and plasmid multilocus sequence typing. Antimicrob. Agents Chemother. 58, 3895–3903 (2014).

42. Seemann, T. Prokka: rapid prokaryotic genome annotation. Bioinformatics 30, 2068–2069 (2014).

43. Florensa, A. F., Kaas, R. S., Clausen, P. T. L. C., Aytan-Aktug, D. & Aarestrup, F. M. ResFinder - an open online resource for identification of antimicrobial resistance genes in next-generation sequencing data and prediction of phenotypes from genotypes. Microb. Genom. 8, 000748 (2022).

44. Feldgarden, M. et al. AMRFinderPlus and the reference gene catalog facilitate examination of the genomic links among antimicrobial resistance, stress response, and virulence. Sci. Rep. 11, 12728 (2021).

45. Siguier, P., Perochon, J., Lestrade, L., Mahillon, J. & Chandler, M. ISfinder: the reference centre for bacterial insertion sequences. Nucleic Acids Res. 34, D32–36 (2006).

46. Ross, K. et al. TnCentral: a prokaryotic transposable element database and web portal for transposon analysis. mBio 12, e0206021 (2021).

47. Néron, B. et al. IntegronFinder 2.0: identification and analysis of integrons across bacteria, with a focus on antibiotic resistance in *Klebsiella*. Microorganisms 10, 700 (2022).

48. Moura, A. et al. INTEGRALL: a database and search engine for integrons, integrases and gene cassettes. Bioinformatics 25, 1096–1098 (2009).

49. Redondo-Salvo, S. et al. COPLA, a taxonomic classifier of plasmids. BMC Bioinformatics 22, 390 (2021).

50. Hancock, S. J. et al. Identification of IncA/C plasmid replication and maintenance genes and development of a plasmid multilocus sequence typing scheme. Antimicrob. Agents Chemother. 61, e01740–16 (2017).

51. Alikhan, N.-F., Petty, N. K., Ben Zakour, N. L. & Beatson, S. A. BLAST Ring Image Generator (BRIG): simple prokaryote genome comparisons. BMC Genomics 12, 402 (2011).

52. Katoh, K. & Standley, D. M. MAFFT multiple sequence alignment software version 7: improvements in performance and usability. Mol. Biol. Evol. 30, 772–780 (2013).

53. Page, A. J. et al. SNP-sites: rapid efficient extraction of SNPs from multi-FASTA alignments. Microb. Genom. 2, e000056 (2016).

54. Stamatakis, A. RAxML version 8: a tool for phylogenetic analysis and post-analysis of large phylogenies. Bioinformatics 30, 1312–1313 (2014).

55. Welch, T. J. et al. Multiple antimicrobial resistance in plague: an emerging public health risk. PLoS ONE 2, e309 (2007).

56. Del Castillo, C. S. et al. Comparative sequence analysis of a multidrug-resistant plasmid from *Aeromonas hydrophila*. Antimicrob. Agents Chemother. 57, 120–129 (2013).

57. Doublet, B. et al. Complete nucleotide sequence of the multidrug resistance IncA/C plasmid pR55 from *Klebsiella pneumoniae* isolated in 1969. J. Antimicrob. Chemother. 67, 2354–2360 (2012).

58. Fricke, W. F. et al. Comparative genomics of the IncA/C multidrug resistance plasmid family. J. Bacteriol. 191, 4750–4757 (2009).

59. Towner, K. J., Pearson, N. J., Mhalu, F. S. & O’Grady, F. Resistance to antimicrobial agents of *Vibrio cholerae* El Tor strains isolated during the fourth cholera epidemic in the United Republic of Tanzania. Bull. World Health Organ. 58, 747–751 (1980).

60. Nigro, S. J. & Hall, R. M. GIsul2, a genomic island carrying the *sul2* sulphonamide resistance gene and the small mobile element CR2 found in the *Enterobacter cloacae s*ubspecies *cloacae* type strain ATCC 13047 from 1890, *Shigella flexneri* ATCC 700930 from 1954 and Acinetobacter baumannii ATCC 17978 from 1951. J. Antimicrob. Chemother. 66, 2175–2176 (2011).

61. Wang, R., Liu, H., Zhao, X., Li, J. & Wan, K. IncA/C plasmids conferring high azithromycin resistance in *Vibrio cholerae*. Int. J. Antimicrob. Agents 51, 140–144 (2018).

62. Zheng, Z. et al. Genetic characterization of *bla*CTX-M-55 -bearing plasmids harbored by food-borne cephalosporin-resistant *Vibrio parahaemolyticus* strains in China. Front. Microbiol. 10, 1338 (2019).

63. Harmer, C. J. & Hall, R. M. IS26 and the IS26 family: versatile resistance gene movers and genome reorganizers. Microbiol. Mol. Biol. Rev. 88, e0011922 (2024).

64. Toleman, M. A., Bennett, P. M. & Walsh, T. R. ISCR elements: novel gene-capturing systems of the 21st century? Microbiol. Mol. Biol. Rev. 70, 296–316 (2006).

65. Partridge, S. R. Analysis of antibiotic resistance regions in Gram-negative bacteria. FEMS Microbiol. Rev. 35, 820–855 (2011).

66. Call, D. R., et al. *blaCMY*-2-positive IncA/C plasmids from *Escherichia coli* and *Salmonella enterica* are a distinct component of a larger lineage of plasmids. Antimicrob. Agents Chemother. 54, 590–596 (2010).

67. Bie, L., Wu, H., Wang, X.-H., Wang, M. & Xu, H. Identification and characterization of new members of the SXT/R391 family of integrative and conjugative elements (ICEs) in *Proteus mirabili*s. Int. J. Antimicrob. Agents 50, 242–246 (2017).

68. Pinilla-Redondo, R. et al. CRISPR-Cas systems are widespread accessory elements across bacterial and archaeal plasmids. Nucleic Acids Res. 50, 4315–4328 (2022).

69. Pinilla-Redondo, R. et al. Type IV CRISPR-Cas systems are highly diverse and involved in competition between plasmids. Nucleic Acids Res. 48, 2000–2012 (2020).

70. Durand, R., Deschênes, F. & Burrus, V. Genomic islands targeting *dusA* in *Vibrio* species are distantly related to *Salmonella* Genomic Island 1 and mobilizable by IncC conjugative plasmids. PLoS Genet. 17, e1009669 (2021).

71. Grim, C. J. et al. Genome sequence of hybrid *Vibrio cholerae* O1 MJ-1236, B-33, and CIRS101 and comparative genomics with V. cholerae. J. Bacteriol. 192, 3524–3533 (2010).

72. Durand, R. et al. Crucial role of *Salmonella* genomic island 1 master activator in the parasitism of IncC plasmids. Nucleic Acids Res. 49, 7807–7824 (2021).

73. Ambrose, S. J. & Hall, R. M. Can SGI1 family integrative mobilizable elements overcome entry exclusion exerted by IncA and IncC plasmids on IncC plasmids? Plasmid 123–124, 102654 (2022).

74. Harmer, C. J., Hamidian, M., Ambrose, S. J. & Hall, R. M. Destabilization of IncA and IncC plasmids by SGI1 and SGI2 type *Salmonella* genomic islands. Plasmid 87–88, 51–57 (2016).

75. Ambrose, S. J. & Hall, R. M. SGI1 excludes IncA and IncC plasmids. Plasmid 133, 102743 (2025).

76. Deschênes, F. et al. Conjugative transfer inhibition of IncA and IncC plasmids by pervasive SGI1-like elements via relaxosome assembly interference. 2025.04.21.649884 Preprint at 10.1101/2025.04.21.649884 (2025).

77. Roy, D., Huguet, K. T., Grenier, F. & Burrus, V. IncC conjugative plasmids and SXT/R391 elements repair double-strand breaks caused by CRISPR-Cas during conjugation. Nucleic Acids Res. 48, 8815–8827 (2020).

78. Nair, S. et al. Highly drug-resistant *Vibrio cholerae* harbouring *blaPER*-7 isolated from travellers returning to England. J. Antimicrob. Chemother. 80, 2428–2432 (2025).

79. Appelt, S. et al. Import of multidrug-resistant *Vibrio cholerae* from Ethiopia to Germany and the UK. Lancet Microbe 6, 101179 (2025).

80. Shaw, S. et al. Genomic portrayal of emerging carbapenem-resistant El Tor variant *Vibrio cholerae* O1. Antimicrob. Agents Chemother. 69, e0074025 (2025).

81. Luo, Y., Payne, M., Kaur, S., Octavia, S. & Lan, R. Genomic evidence of two-staged transmission of the early seventh cholera pandemic. Nat. Commun. 15, 8504 (2024).

82. Folster, J. P. et al. Multidrug-resistant IncA/C plasmid in *Vibrio cholerae* from Haiti. Emerg. Infect. Dis. 20, 1951–1953 (2014).

83. Wang, R., Li, J. & Kan, B. Sequences of a co-existing SXT element, a chromosomal integron (CI) and an IncA/C plasmid and their roles in multidrug resistance in a *Vibrio cholerae* O1 El Tor strain. Int. J. Antimicrob. Agents 48, 305–309 (2016).

