## Supplementary Figures for "Evolution of multidrug-resistant IncC plasmids in the seventh pandemic *Vibrio cholerae* O1 El Tor lineage between 1979 and 2024"

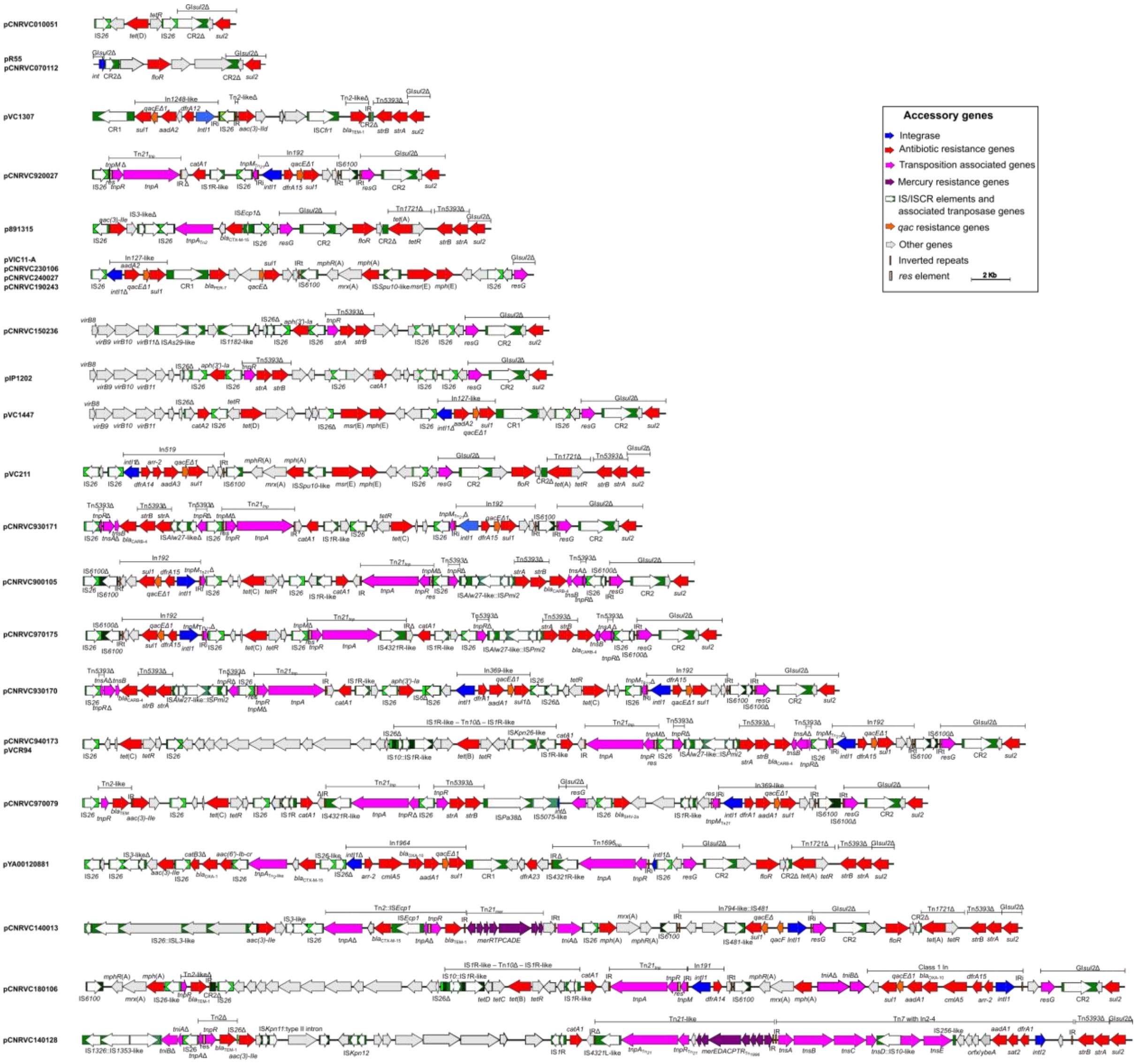


**Supplementary Figure 1: ARI-B structures in IncC2 plasmids.** Arrows indicate gene length and orientation, with gene names shown above or below the arrows and colors corresponding to the associated key. The boundaries of retrieved modules are shown above the genomic region. Boxes in different shades of green represent insertion sequences (ISs) and common regions (CRs), with white arrows indicating the direction of transposition-associated genes. Class 1 integrons were identified based on their cassette arrays as annotated in the Integrall database. Complex integrons were inferred based on the presence of inverted repeats (IRi and IRt**)**. The scale bar indicates the size of accessory structures.

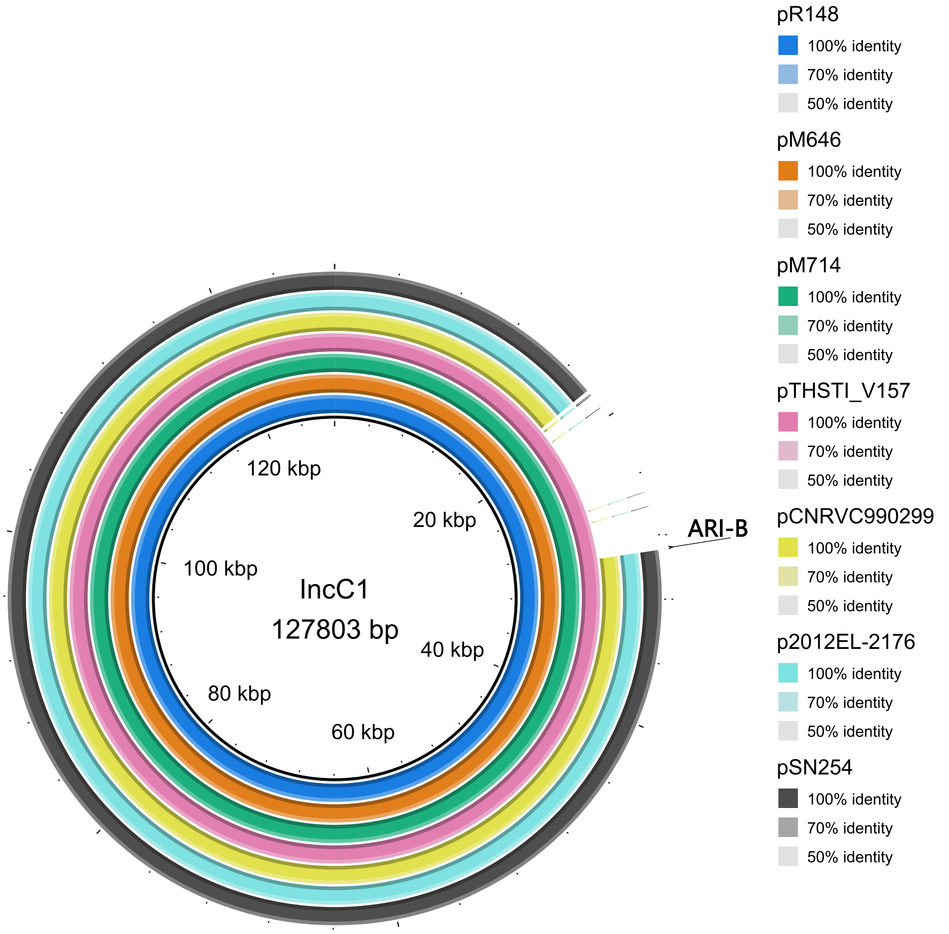


**Supplementary Figure 2. Alignment of seven IncC1 plasmids against the IncC1 backbone.** The innermost circle shows the nucleotide positions of the IncC1 plasmid backbone derived from pR148 (GenBank accession no. JX141473.1). The subsequent colored circles show the alignments of seven plasmids with the IncC1 backbone. The arrow shows the location of ARI-B. A backbone region adjacent to the ARI-B insertion site is deleted in pCNRVC990299, p2012EL-2176, and pSN254.

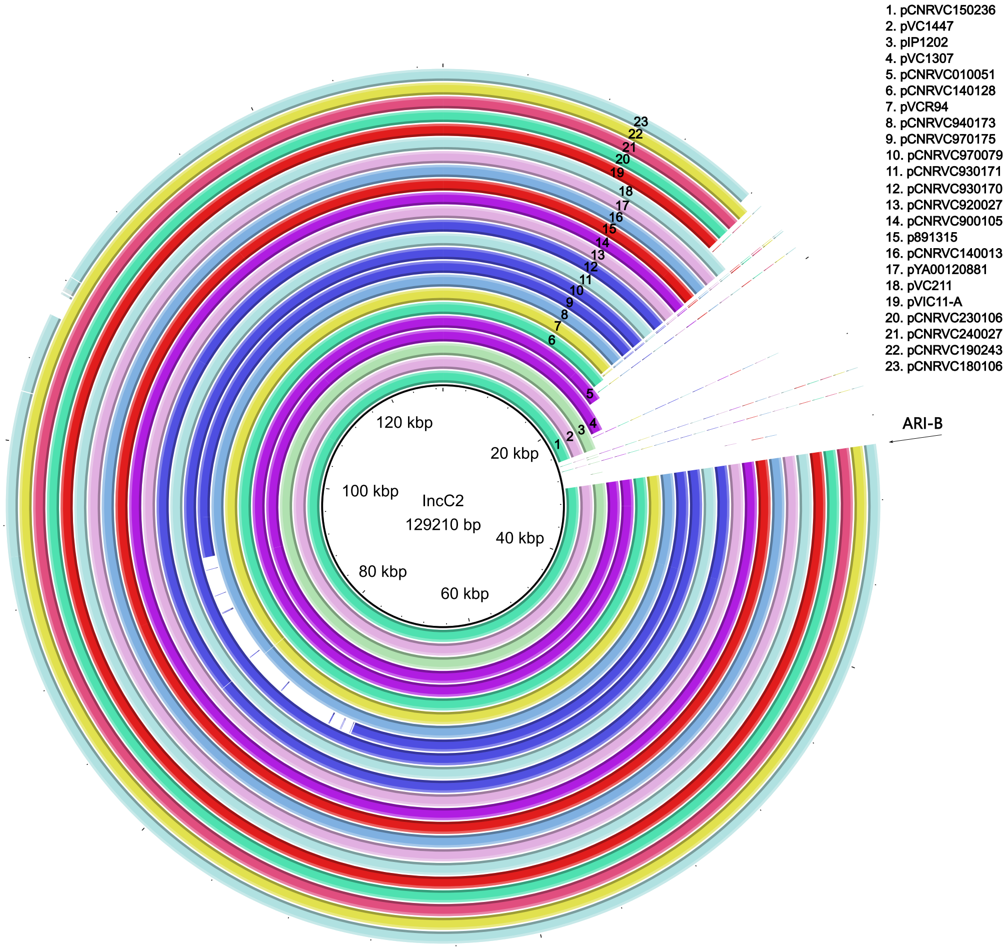


**Supplementary Figure 3. Alignment of 23 IncC2 plasmids with backbone deletions adjacent to the ARI-B insertion site against the IncC2 backbone.** The innermost circle indicates the nucleotide positions of the IncC2 plasmid backbone derived from pR55 (GenBank accession no. JQ010984.1); circles 1–23 show alignments of the individual plasmids with the IncC2 backbone. The arrow shows the location of ARI-B.

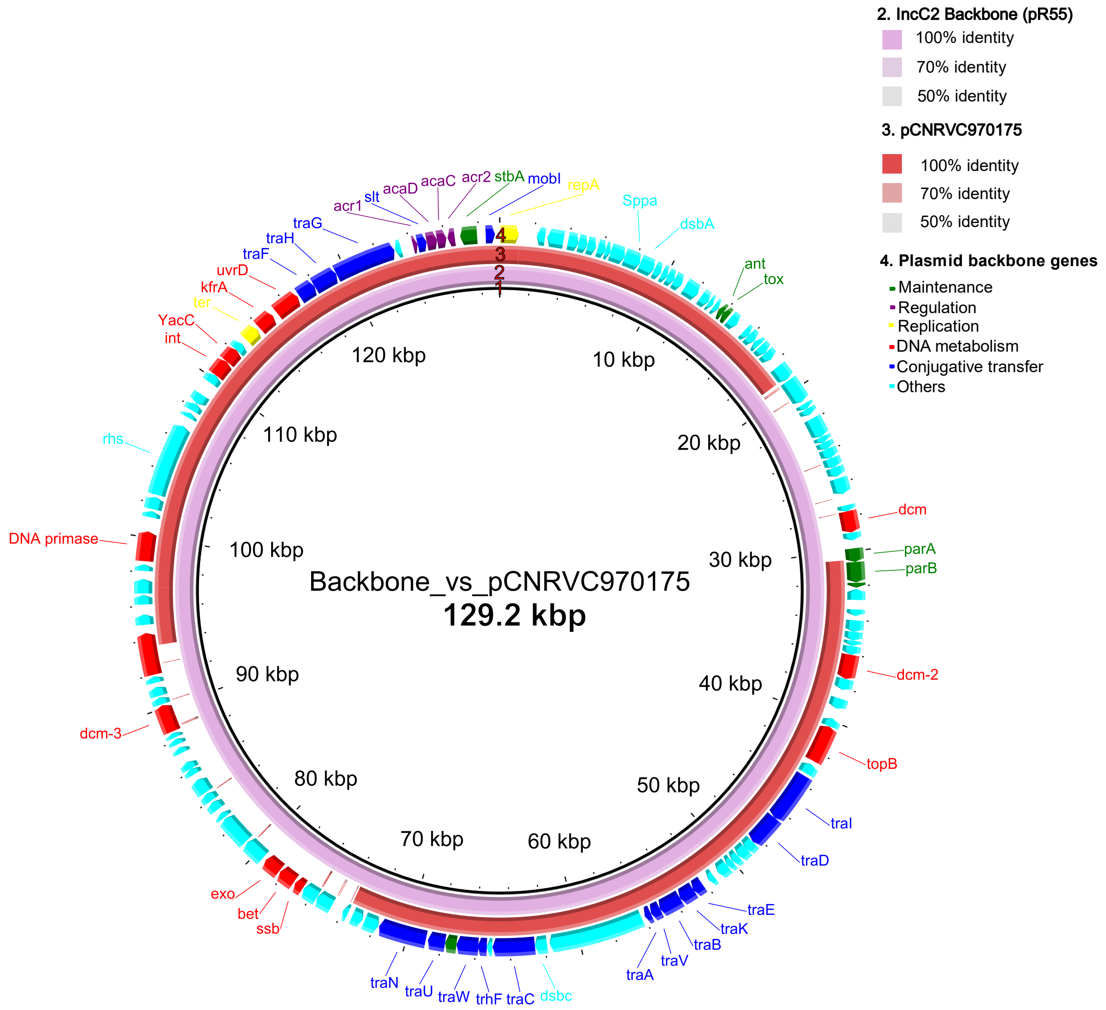


**Supplementary Figure 4. Circular map and comparative analysis of the IncC2 backbone and pCNRVC970175.** The innermost circle (circle 1) indicates the nucleotide positions of the IncC2 plasmid backbone derived from pR55 (GenBank accession no. JQ010984.1). Circles 2 and 3 show the alignments of pR55 and pCNRVC970175, with colors indicating the percentage of nucleotide identity. Arrows indicate the names of selected genes in circle 4.
